# Community composition and the environment modulate the population dynamics of type VI secretion in human gut bacteria

**DOI:** 10.1101/2023.02.20.529031

**Authors:** Sophie Robitaille, Emilia L. Simmons, Adrian J. Verster, Emily Ann McClure, Darlene B. Royce, Evan Trus, Kerry Swartz, Daniel Schultz, Carey D. Nadell, Benjamin D. Ross

## Abstract

Understanding the relationship between the composition of the human gut microbiota and the ecological forces shaping it is of high importance as progress towards therapeutic modulation of the microbiota advances. However, given the inaccessibility of the gastrointestinal tract, our knowledge of the biogeographical and ecological relationships between physically interacting taxa has been limited to date. It has been suggested that interbacterial antagonism plays an important role in gut community dynamics, but in practice the conditions under which antagonistic behavior is favored or disfavored by selection in the gut environment are not well known. Here, using phylogenomics of bacterial isolate genomes and analysis of infant and adult fecal metagenomes, we show that the contact-dependent type VI secretion system (T6SS) is repeatedly lost from the genomes of *Bacteroides fragilis* in adults compare to infants. Although this result implies a significant fitness cost to the T6SS, but we could not identify *in vitro* conditions under which such a cost manifests. Strikingly, however, experiments in mice illustrated that the *B. fragilis* T6SS can be favored or disfavored in the gut environment, depending on the strains and species in the surrounding community and their susceptibility to T6SS antagonism. We use a variety of ecological modeling techniques to explore the possible local community structuring conditions that could underlie the results of our larger scale phylogenomic and mouse gut experimental approaches. The models illustrate robustly that the pattern of local community structuring in space can modulate the extent of interactions between T6SS-producing, sensitive, and resistant bacteria, which in turn control the balance of fitness costs and benefits of performing contact-dependent antagonistic behavior. Taken together, our genomic analyses, *in vivo* studies, and ecological theory point toward new integrative models for interrogating the evolutionary dynamics of type VI secretion and other predominant modes of antagonistic interaction in diverse microbiomes.

## INTRODUCTION

The human gastrointestinal tract harbors a dense microbial community, collectively referred to as the gut microbiota, that influences numerous aspects of human physiology including immune development, metabolism, and the gut-brain axis, among others ^1–3^. Across human populations, the composition of the gut microbiota exhibits striking diversity ^4–6^. How these patterns arise and what forces shape microbiota assembly and stability are not fully understood; filling this knowledge gap is essential for our fundamental understanding of microbial ecology and for the future improvement of microbiome-directed therapies ^7^. While extrinsic factors such as host genetics and diet can play key roles in influencing the gut microbiota, the importance of interbacterial interactions in shaping gut microbiome composition, stability, and function is an emerging area of increasing research focus ^8,9^. To-date*, in situ* interbacterial interactions in the human gut microbiota have largely been inferred indirectly via association studies of large metagenomic datasets^10^. Direct evidence of relevant physical associations between discrete taxa has been impeded by the difficulty of gaining physical access to the unperturbed healthy human intestine. Studies in mice have revealed that gut bacterial biogeography is not homogeneous but is instead highly structured at the micrometer scale, with evidence for biofilm-like assemblages whose composition depends on longitudinal localization in the intestine ^11^. New technology may yield some insight into the physiology and localization of gut microbes, however aside from limited cases, it is likely that a combination of different approaches is necessary to infer how localized microbiota structuring might influence community assembly and dynamics through time ^9^.

Direct antagonistic interactions are thought to be pervasive in the gastrointestinal tract, given the diversity and prevalence of pathways encoding attack mechanisms that are carried by gut symbionts ^12,13^. One well-studied antagonistic pathway is the type VI secretion system (T6SS) encoded by Gram-negative bacteria, including the highly abundant gut taxon Bacteroidales and many enteric pathogens ^14^. The T6SS is a contact-dependent protein translocation system used to deliver bacteriostatic or bacteriolytic effector toxins into target cells ^15,16^, thereby potentially affecting microbiota composition through the elimination of susceptible bacteria and opening of space and resources for T6SS-producing and resistant cells. The role of the T6SS in mammalian gut microbiome ecology is an emerging field. T6SS activity during invasion by enteric pathogens, including species of the *Shigella*, *Salmonella*, and *Citrobacter* genera, facilitates colonization and subsequent pathogenesis typically through the eradication of phylogenetically related endogenous symbionts ^17–19^. In the human gut, the most abundant T6SS-encoding taxon is the order Bacteroidales, including species of the *Parabacteroides* and *Bacteroides* genera ^20^. The T6SS of these organisms mediates inter-species and inter-strain competition during experimental colonization of mice and is inferred by metagenomic studies to be an important factor for their presence in the human gut ^21–25^.

Despite its demonstrated utility and impact, the T6SS is not ubiquitous in all human gut microbiomes, and some Bacteroidales genomes possess mutationally inactivated T6SS pathways; others lack the T6SS altogether ^22,24–26^. These paradoxical observations suggest the possibility that the competitive benefits of T6SS activity can be offset by costs that manifest in certain gut environments or ecological contexts, leading to elimination of T6SS function. To-date this trade-off has not been experimentally explored. Here, we examine the population and evolutionary dynamics of T6SS-producing and non-producing variants of the human gut symbiont *Bacteroides fragili*s through metagenomic, experimental, and theoretical approaches. We document numerous instances from human microbiome data recapitulating the evolutionary loss of T6SS, and we explore experimentally how the population dynamics of T6SS secretion depend on composition and T6SS susceptibility of the surrounding community. Finally, we use two different modeling approaches to interrogate potential microenvironmental conditions that could explain variance in T6SS population and evolutionary dynamics as a function of community composition. Our work underscores that the importance of antagonistic interbacterial interactions in the human gut microbiome depends on context, providing mechanistic insight into the selective pressures that contribute to differences in microbiome composition. Further exploration of interbacterial interactions in the gut may lead to deeper insight into their ecological relevance and evolutionary dynamics, better informing future efforts to manipulate these communities.

## RESULTS

### The *Bacteroides fragilis* GA3 T6SS is enriched in infants and associated with specific *Bacteroides* species in adults

Bacteroidales species utilize the T6SS to compete with related bacteria within the mammalian gut^23^. Bacteroidales-encoded T6SS comprise three distinct subtypes, termed genetic architecture (GA) 1-3 ^27^. While GA1 and GA2 are harbored on mobile integrative and conjugative elements (ICE) that transfer between co-resident species, the GA3 system is restricted to *Bacteroides fragilis.* We previously demonstrated that *B. fragilis* strains residing within the adult gut microbiome often lack the T6SS GA3 subtype, and that the GA3 T6SS is more prevalent in infant microbiomes ^24^. We hypothesized that these observations reflect selection for loss of the GA3 T6SS over the course of within-person evolution. To further investigate this hypothesis, we first reanalyzed a large number of infant and adult microbiome samples containing *B. fragilis* to gain more detailed insight into patterns of GA3 presence and absence ^28,29^. We found that, regardless of host age, *B. fragilis* abundance largely exhibited a strong positive correlation with the abundance of the GA3 T6SS (Fig. 1a-b). However, for 8% of infant (DIABIMMUNE, N=416) and 17% of adult (HMP, N=194) samples containing *B. fragilis,* GA3 T6SS genes were either not detected or detected at abundance levels lower than those of *B. fragilis,* replicating previous findings in a larger cohort. The difference in the prevalence of GA3-negative *B. fragilis* strains between datasets suggests that independent T6SS loss events occur in the microbiomes of some individuals over the course of human lifespan.

**Fig. 1.**
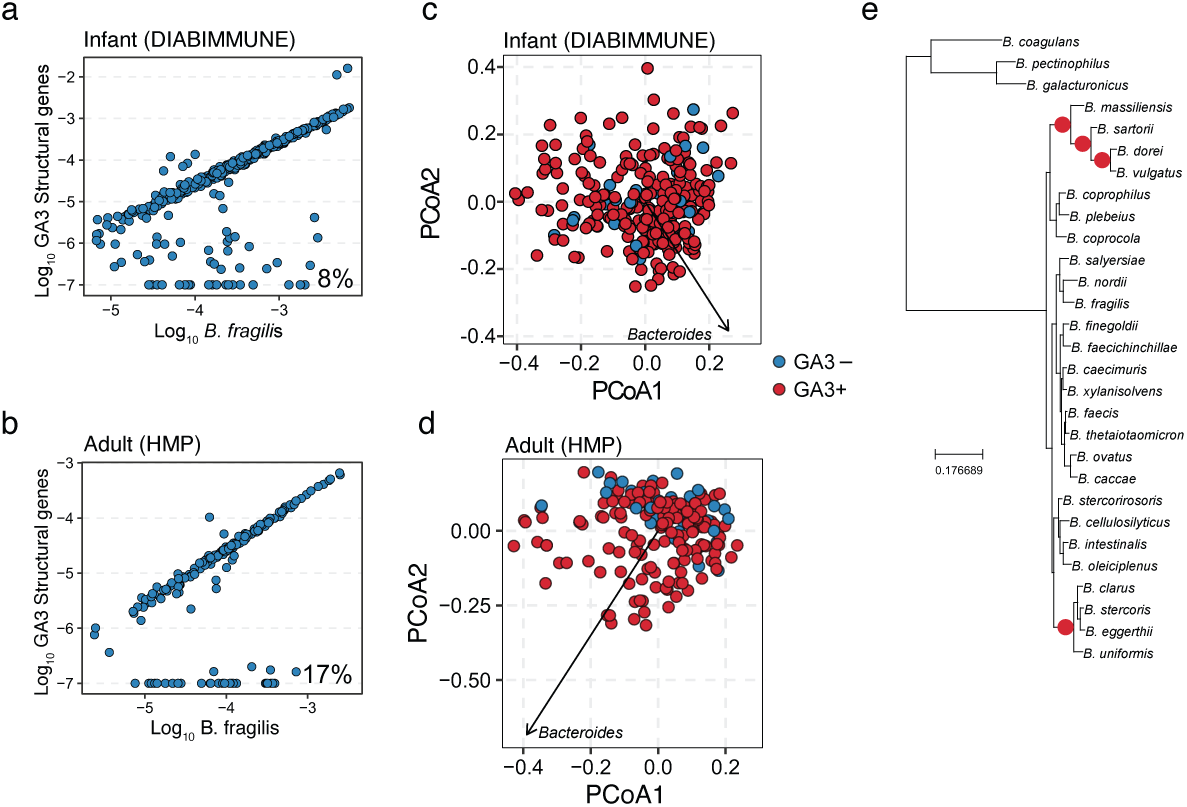
The GA3 T6SS is associated with altered composition of the human gut microbiota. Comparison of abundance of *B. fragilis*-specific GA3 structural genes with *B. fragilis* marker genes in microbiome samples from (**a**) 6-to-18 month-old infants (DIABIMMUNE, N=416) and (**b**) adults (Human Microbiome Project, N=440). Scatter plots depict the relationship between the log10 abundance of *B. fragilis* and the log10 abundance of GA3 for samples in which at least 100 reads map to *B. fragilis* marker genes. GA3 abundance in each sample was adjusted with a pseudo count of 1e-7 and thus samples without detectable GA3 are present at -7. The percentage of *B. fragilis-*containing samples lacking GA3 is indicated at lower right in each plot. (**c-d**) PcoA plots depicting *B. fragilis*-positive shotgun metagenomic samples from (**c**) 6-to-18 month old infants from the DIABIMMUNE cohort and (**d**) adult HMP cohort. Samples are colored by the presence (red) or absence (blue) of the GA3 T6SS. Distances between samples were calculated using unweighted unifrac with genus-level relative abundance values and a phylogenetic tree of bacterial genera. A single loading vector is shown for *Bacteroides* and the separation between GA3+ and GA3- was assessed in each plot using PERMANOVA tests (DIABIMMUNE p > 0.05, HMP p < 0.01). **©** A phylogenetic tree of *Bacteroides* species that highlights significant PhILR nodes in red. We used the PhILR algorithm with Bonferroni correction for multiple tests to identify significant nodes on the MetaPhlAn3 phylogenetic tree using species relative abundance values.

To explore differences in community composition between microbiome samples in which *B. fragilis* possessed or lacked the GA3 T6SS, we performed principal components analysis of the genus-level taxonomic profiles of each sample. Strikingly, while infant microbiome samples harboring *B. fragilis* could not be distinguished based on the presence of the T6SS (Fig. 1c, PERMANOVA p > 0.05), we found that adult microbiomes exhibited significant separation between T6SS+ and T6SS- samples (Fig. 1d, PERMANOVA, p < 0.01) along an axis of *Bacteroides* genus relative abundance, consistent with previous findings ^24^. To investigate if this separation was driven by individual *Bacteroides* species, we used the phylogenetic log ratio transform algorithm to compare species relative abundances between T6SS+ and T6SS- adult microbiome samples ^30^. This analysis revealed that the *B. fragilis* T6SS+ was associated with the increased relative abundance of two distinct clades of *Bacteroides* species, including the highly abundant and prevalent species *B. vulgatus* and *B. uniformis* (Fig. 1e, Supplementary Table 1). Importantly, the phylogenetic targeting breadth of the *B. fragilis* GA3 T6SS encompasses these clades, and previous results have shown that these species are likely to be susceptible to GA3 T6SS intoxication in the gut ^21^. In sum, these findings demonstrate that the *B. fragilis* GA3 T6SS is likely to be lost over time in the human gut, and its presence in adults is associated with the relative abundance of specific susceptible target species, an association which likely develops over the course of human lifespan.

### Parallel loss of the *B. fragilis* GA3 T6SS

Parallel evolutionary events of GA3 loss across the *B. fragilis* phylogeny suggest selection against retention of this trait in some contexts, and, since *B. fragilis* strains largely exist as clonal populations in each microbiome ^24,29,31^, we reasoned that these patterns provided support for our previous inference that the GA3 system can be selected against over human developmental time ^24^. To further investigate this idea, we searched 387 *B. fragilis* genomes for the presence of 13 GA3 core genes and performed phylogenetic analysis. We found that 288 of the 387 (74.4%) genomes possessed all GA3 core genes, while 99 (25.6%) genomes lacked all or some of these genes (Supplementary Table 2). Upon close examination of the syntenic region of the *B. fragilis* chromosome harboring the GA3 T6SS locus in a subset of genomes possessing or lacking the system, we found evidence of varied levels of genomic erosion including point mutations introducing premature stop codons, partial loss of a subset of T6SS genes encoding structural components of the apparatus, loss of the entire GA3 system, and even loss of the greater locus including unrelated syntenic flanking genes (Fig. 2a, Supplementary Table 2). We next used ancestral state reconstruction to infer the evolutionary relationships between loss events across the phylogeny, detecting 18 independent events (Fig. 2b). Noticing that most inferred loss events occurred at the tips of the *B. fragilis* phylogenetic tree instead of at deeper nodes, we calculated the distance-to-tip for each inferred loss event and compared it to the mean distance for 1000 simulated trees in which phylogenetic relationships were randomized. While the observed median distance to tip was 0.00052, the median across all simulations was 0.00598 and was greater than the observed distance in every simulated tree (empirical P < 0.001). To confirm that our results were robust to phylogenetic tree construction, we repeated our analysis over 1000 bootstrap trees and found a significant (P < 0.05) result in 98.1% of trees (median empirical P = 0.001). We note that there is one clade that has lost GA3 in our phylogenetic tree, however, our analysis suggests this is an outlier and the usual result of GA3 loss is rapid extinction. This finding suggests that the GA3 T6SS tends to be retained across the *B. fragilis* phylogeny; genomic loss of GA3, therefore, appears often to be transient over human lifespan within individuals, but GA3-containing lineages may persist more often over longer time scales involving transfer from one host to another. While the T6SS may be common and advantageous for *B. fragilis* during early human gut colonization, a sizable fraction of host microbiome conditions occurring across the human lifespan lead to the parallel evolutionary loss of the T6SS ^31,32^.

**Fig. 2.**
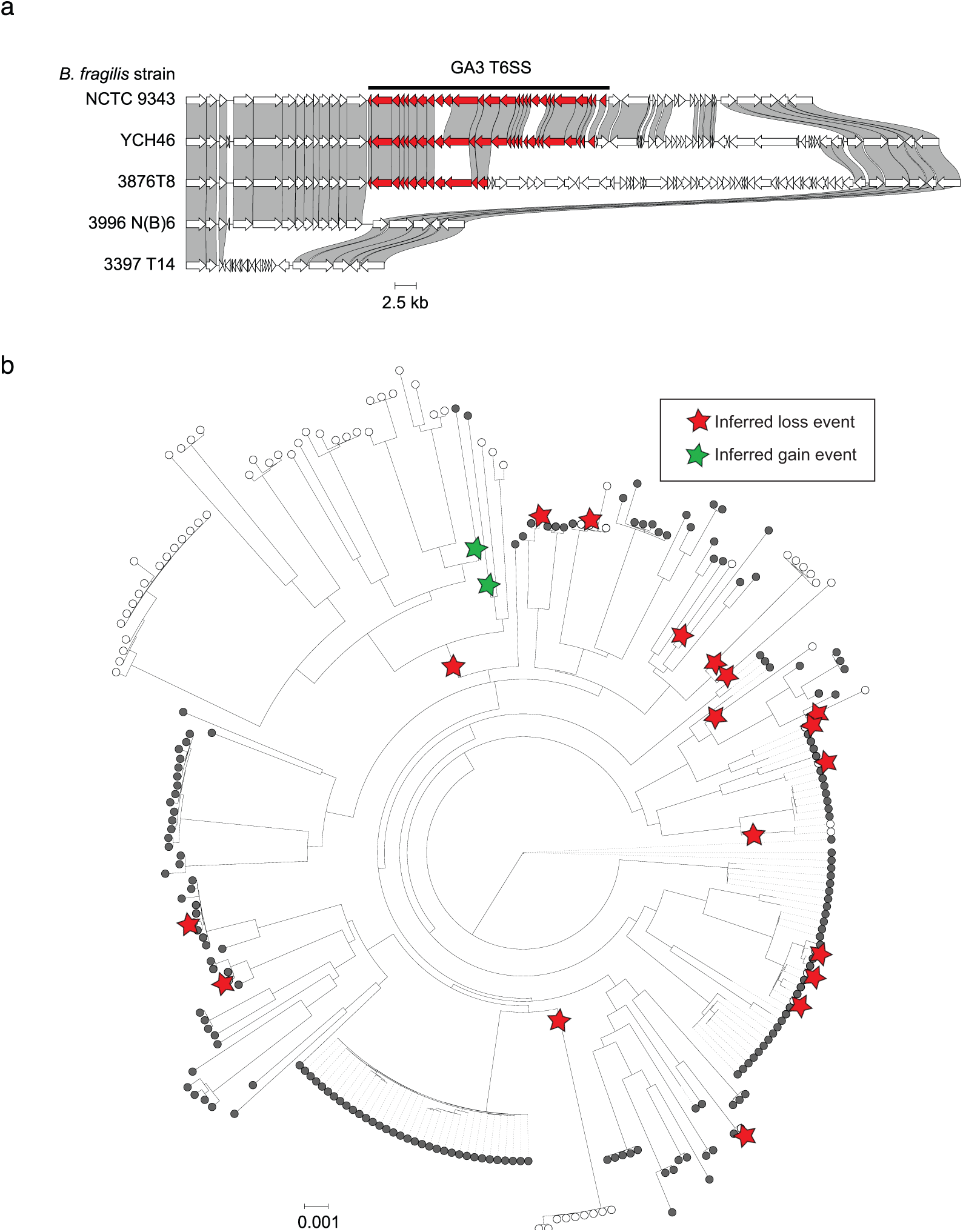
Recurrent evolutionary loss of the GA3 T6SS from *B. fragilis.* (**a**) Schematic comparing intact and degenerated GA3 loci (red genes) from syntenic regions of select *B. fragilis* genomes. Homology is indicated by shaded regions linking genes between genomes of different strains. (**b**) Maximum likelihood phylogenetic tree of *B. fragilis* strains based on single nucleotide polymorphisms in aligned *B. fragilis* MetaPhlAn3 marker genes. GA3 presence and absence are indicated by closed and open circles (respectively) at branch tips. Inferred unique loss (red) and gain (green) events during *B. fragilis* diversification are indicated by stars, based on ancestral state reconstruction.

### Loss of T6SS function does not confer an advantage during growth *in vitro*

Based on our initial results, we hypothesized that the GA3 T6SS confers a fitness cost to *B. fragilis* strains, resulting in selection for strains that have inactivated the pathway in some contexts. To begin to test this idea, we performed liquid growth assays comparing a parental *B. fragilis* “Producer” strain and a “Resistant” strain lacking a functional GA3 pathway through in-frame deletion of the sheath genes *tssC-B* (BF9343_1941-2) and the ATPase gene *clpV* (BF9343_1940) but retaining immunity genes (Fig. S1). Experiments were performed in defined minimal media with a variety of different carbon sources, including mono-, di-, and polysaccharides (Extended Data Fig. 1-2). We hypothesized that in media with carbon sources such as inulin or D-xylose, on which *B. fragilis* has a low growth rate ^33^, any growth rate cost of T6SS expression and activity might be observed in a growth curve. However, we observed no detectable difference in growth rate or maximal optical density between *B. fragilis* strains in any liquid media condition (Extended Fig. 1-2). Similar results were found for a “Resistant” strain lacking only *tssC* (Fig. S2). Although *B. fragilis* strains secrete the T6SS needle tube protein Hcp under growth in planktonic conditions ^34^, it is well-established that the function of the T6SS in interbacterial antagonism depends on prolonged cell-cell contact such as during surface-associated growth or in highly viscous environments ^16,35^. We therefore hypothesized that liquid conditions might not trigger full activation of the system, masking any potential fitness differences between Producer and Resistant strains. However, mono- and co-culture assays on defined minimal media agar plates with the same sole carbon sources and on mucin also failed to reveal any growth advantage of the resistant strain (Extended Data Fig. 1-2). In sum, our results indicated that the presence of an intact T6SS pathway does not significantly impact the growth rate of *B. fragilis* under *in vitro* conditions.

### A substantial cost of T6SS-production manifests in the mouse intestinal environment

Growth *in vitro* does not recapitulate *in vivo* population sizes, cellular physiology, growth dynamics, or spatial population structure in the digestive tract, all of which could influence the impact of T6SS activity on *B. fragilis* fitness. In both antibiotic-treated conventionally-reared mice and gnotobiotic mice, the activity of the GA3 T6SS dramatically influences the outcome of competition between *B. fragilis* strains during co-colonization by reducing the abundance of susceptible cells ^21,23,25^. We therefore sought to examine the impact of the T6SS on *B. fragilis* fitness during colonization of mice. We gavaged antibiotic-treated specific pathogen free mice with a 1:1 mixture of *B. fragilis* Producer and Resistant strains and tracked the relative abundance of these strains in fecal pellets collected over the course of approximately six months (Fig. 3a). We also performed 16S rRNA gene amplicon sequencing and profiled taxonomic relative abundance of the residual endogenous microbiota over time. As expected, at early timepoints, the dominant genus was *Bacteroides* (Fig. S3). However, at later timepoints in the majority of mice, the endogenous microbiota increased in relative abundance, including several non-*B. fragilis* taxa from the order Bacteroidales. Notably, we found that Producer strain abundance rapidly and reproducibly decreased relative to the Resistant strain, such that by the final timepoint the average Producer percentage in pellets was 12% of the sum of Producer and Resistant gDNA detected, with multiple mice harboring less than 0.5% (Fig. 3b-c). Similar percentages of Producer were detected in the small intestine, cecum, and colon following sacrifice (Fig. S4). These findings mirrored parallel mice colonization studies with a Resistant strain bearing only a single inactivating mutation (Δ*tssC*) (Fig. S5). In a minority of mice, at later time points, we observed a partial rebound in Producer frequency; this was not associated with any consistent patterns in microbiome composition relative to mice in which Producer abundance remained low. Further functional analysis of isolates from mice in which the Producer rebound occurred gave no indication of T6SS inactivation (i.e. conversion of Producer mutants to Resistant cells) (Fig. S6), and indeed we detected no mutations in or near the GA3 T6SS operon upon whole genome sequencing (Supplementary table 3). The basis for Producer rebound thus was not entirely resolved.

**Fig. 3.**
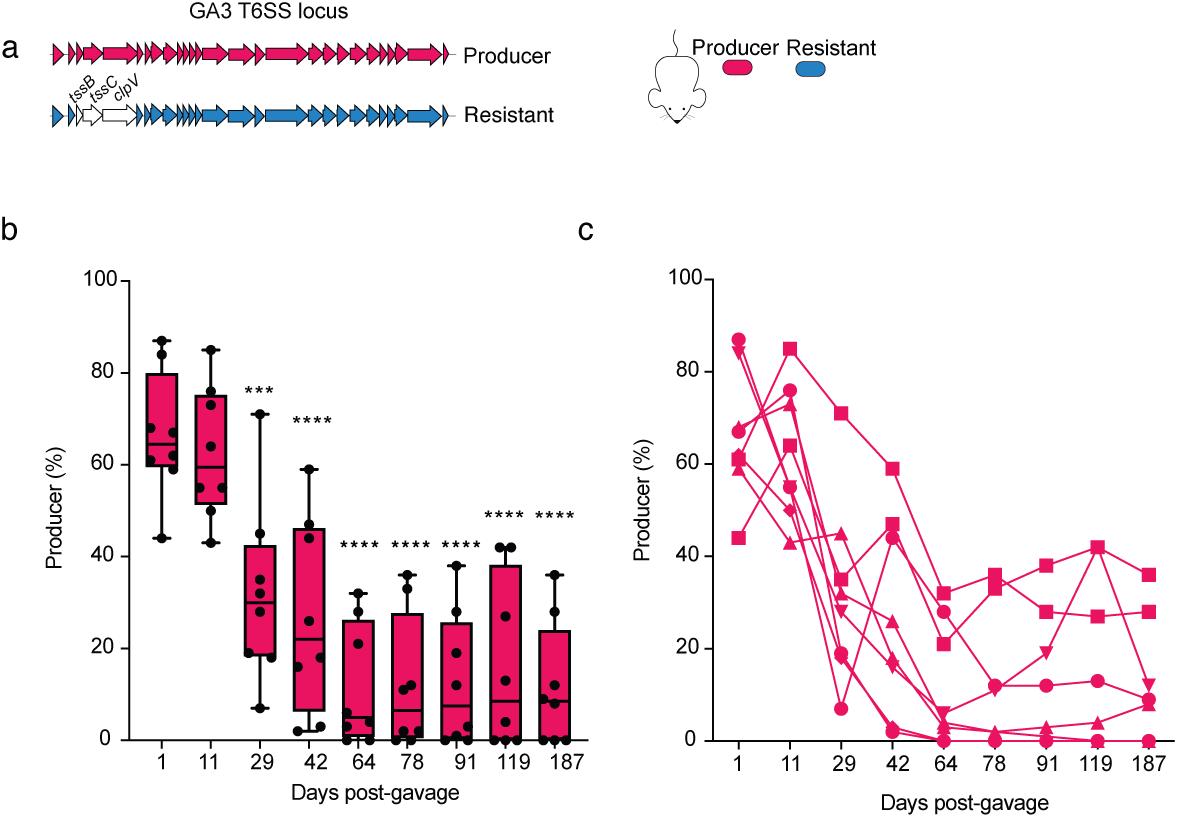
A Resistant strain outcompetes a Producer strain during co-colonization of mice. (**a**) Schematic representing the *B. fragilis* NCTC 9343 GA3 T6SS operon, comparing the “Producer” T6SS- active strain (magenta) to the “Resistant” T6SS-inactive strain (blue). The in-frame 3-gene deletion mutation (Δ*tssBC-clpV*) in the resistant strain is indicated by lack of shading. Strains were inoculated via oral gavage of a 1:1 mixture into antibiotic-treated SPF mice. Strain relative abundance (normalized to 100%) was tracked via barcode-specific quantitative PCR performed on DNA isolated from fecal pellets at indicated days post gavage. (**b**) Quantification of the percent relative abundance of Producer strain compared to the total quantity of gDNA detect for both Producer and Resistant strains (n=8 mice). Box plots indicate the mean and interquartile range. One-way ANOVA test was used to compare abundance between sampling at day 1 (one day after gavage) and all subsequent sampling (***P<0.001, ****P<0.00001). (**c**) Percent abundance of producer strain in each of 8 mice in compared to the total quantity of gDNA detect for both Producer and Resistant strains (**b**).

The results above demonstrate that in a more realistic mouse gut environmental setting, by contrast with our *in vitro* experiments, we can discern a consistent and considerable cost to T6SS production. We note though that in these experiments, the Producer strain is only introduced with a strain against which its T6SS is not effective. The benefit of T6SS, namely killing of competitors to make available additional space and resources, cannot manifest in the absence of competitors that are susceptible to T6SS. In the natural context of the human gut microbiome, however, susceptible competitors are commonly found co-resident with T6SS-encoding strains of *B. fragilis* (Fig. 1a-d).

### The benefit of T6SS production tracks with the presence of sensitive target cells in the mouse gut

To explore how the introduction of a Sensitive strain to our experimental system might alter the population dynamics of T6SS production, we performed a new set of mouse inoculation experiments like those above, but with a 1:1:1 introduction of the Producer strain, the Resistant strain, and a Sensitive strain lacking a functional T6SS and both effector–immunity (E–I) pairs (Fig. 4a and S1). In contrast with the dual colonization experiments, the Producer strain relative abundance strongly increased following gavage, with maximum abundance on day 7. This increase in T6SS producer frequency corresponded with a decrease in the relative abundance of the Sensitive strain (Fig. 4b-g). After this initial advantage to the Producer, however, we observed a return to the pattern observed previously: subsequent sampling revealed a rapid and significant decrease in abundance of the Producer strain to a final mean abundance of 24% with, as before, no difference in percentage between strains in different regions of the intestine (Fig. S7), similar to the outcome observed in the Producer-Resistant two-strain inoculation experiment (Fig. 3b-c, Fig. S4) at day 80, with no mice showing full displacement of the Producer cells. This may reflect the initial boost in abundance the T6SS Producer strain obtained while Susceptible cells were present.

**Fig. 4.**
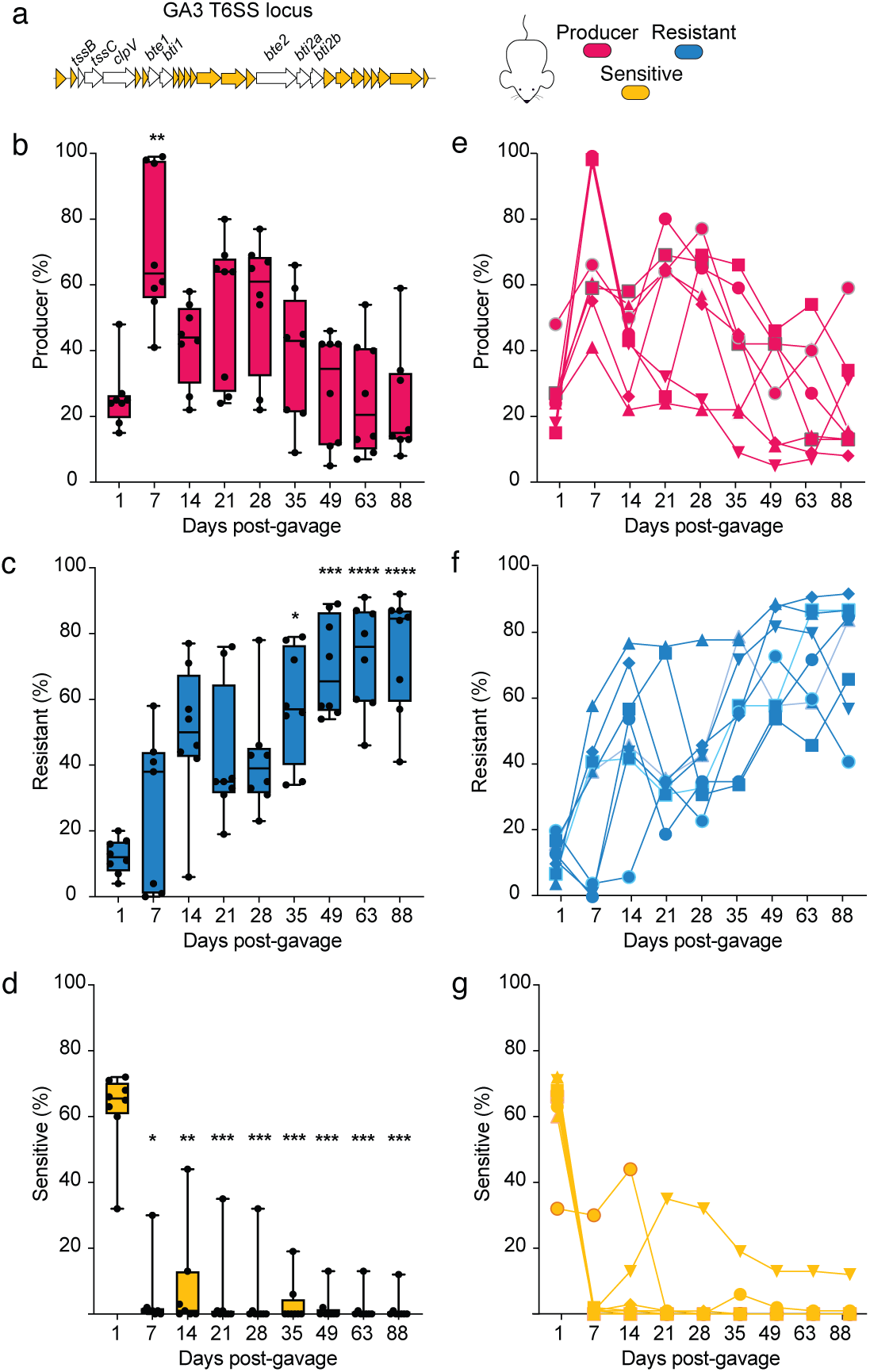
Co-colonization with a Sensitive strain alters Producer-Resistant strain dynamics in mice. (**a**) Schematic representing the *B. fragilis* NCTC 9343 GA3 T6SS operon from the Sensitive strain. The in-frame deletion mutations (Δ*E-I*Δ*tssBC-clpV*) are indicated by lack of shading. Producer, Resistant, and Sensitive strains were inoculated in a 1:1:1 ratio and the gDNA quantity of each strain was tracked for 88 days via qPCR on genomic DNA isolated from fecal pellets. Percentages of Producer (**b**), Resistant (**c**), and Sensitive (**d**) strains are shown as the mean of eight mice for each sampling from days 1 to 88. Significance between mean detected abundance for each strain comparing between day 1 after gavage and all subsequent samplings was assessed by Kruskal-Wallis test (*P<0.05 **P<0.01 ***P<0.001 ****P<0.0001). The percentage of detected Producer (**e**), Resistant (**f**), and Sensitive (**g**) gDNA is shown for each mouse individually over time. Results are representative of two independent colonization experiments, each with N = 8.

### Modeling T6SS competition reveals importance of spatial constraint on population dynamics

Our mouse model experiments provided sufficient environmental realism for us to document that T6SS secretion does indeed carry a cost, and that this cost leads to a repeatable reduction in relative abundance of Producers in the presence of a Resistant strain. When a Sensitive strain is present, killing by the Producer cells appears to offset the cost paid for this behavior, until the Sensitive population is effectively eliminated and the system returns to the Producer-Resistant population dynamics. Recalling the metagenomic data that originally motivated these experiments (Fig. 1), however, there is a substantial fraction of human adults in which Producer and Resistant variants of *B. fragilis* either failed to ever invade or were displaced by Sensitive strains lacking GA3. Though our mouse experiments captured Resistant strains faring better than Producing cells in competition, we did not document Sensitive cells displacing them, a process that might happen at longer timescales than allowed by our experimental setup. To explore possibilities for how these different population dynamic patterns might emerge depending on growth conditions, we turned next to mathematical modeling of the dynamics of these different subpopulations.

To follow the populations of the Producer, Resistant and Sensitive genotypes over time, we model their population dynamics with coupled differential equations of logistic growth, reflecting competition for resources between the different genotypes, with fitness costs for resistance and production of T6SS. We also include a term for contact killing of the Sensitive genotype by the Producer (see Methods) ^36^. After appropriately scaling the variables to arrive at non-dimensional equations, we find that the behavior of our system depends only on only two parameters: the relative costs of T6SS production and resistance, and a scaled “density” parameter *g* that relates the contact killing rate to the carrying capacity, which measures the population density supported by the environment. Scaling of the killing rate by the carrying capacity occurs because interactions between individuals have two distinct negative effects: resource competition and contact inhibition. When the carrying capacity is low, interactions are dominated by resource competition, and contact inhibition becomes less relevant. In environments with a high carrying capacity, supporting dense populations, competition for resources is less intense, increasing the relative importance of contact inhibition. This scaling suggests a general inference that high density of microbial communities in the gut is essential for the fitness advantages provided by production of T6SS.

We then simulated the dynamics of mixed populations for various parameters and found that the fitness of each genotype depends strongly on the density parameter *g*. In low density systems, where contact killing is less relevant, the Sensitive genotype easily dominates the competition due to its higher growth rate (Fig. 5a-f). At higher densities, the Producer genotype does well initially by killing sensitive cells. However, when the elimination of sensitive cells significantly decreases the advantage of T6SS production, the Producer genotype is replaced by the Resistant one, a finding reminiscent of our mouse experiments (Fig. 4). Finally, the dominant genotype at the end of the competition depends on if the elimination of Sensitive cells by the producer was partial or complete. If a subpopulation of Sensitive cells still exists after the Resistant genotype replaces the Producer, the Sensitive genotype can then take advantage of the absence of contact killing and dominate the resistant Strain. Such a scenario is more typical of moderately dense populations, where the contact killing of Sensitive cells is less pronounced. Otherwise, if Sensitive cells are completely killed by the Producer in the beginning of colonization, the Resistant phenotype remains dominant. Notably, complex behaviors with alternation of the dominant genotype are obtained even with relatively small fitness costs for production and resistance (3% and 2% here), which may explain the difficulty of detecting fitness costs of T6SS production in our *in vitro* experiments.

**Fig. 5.**
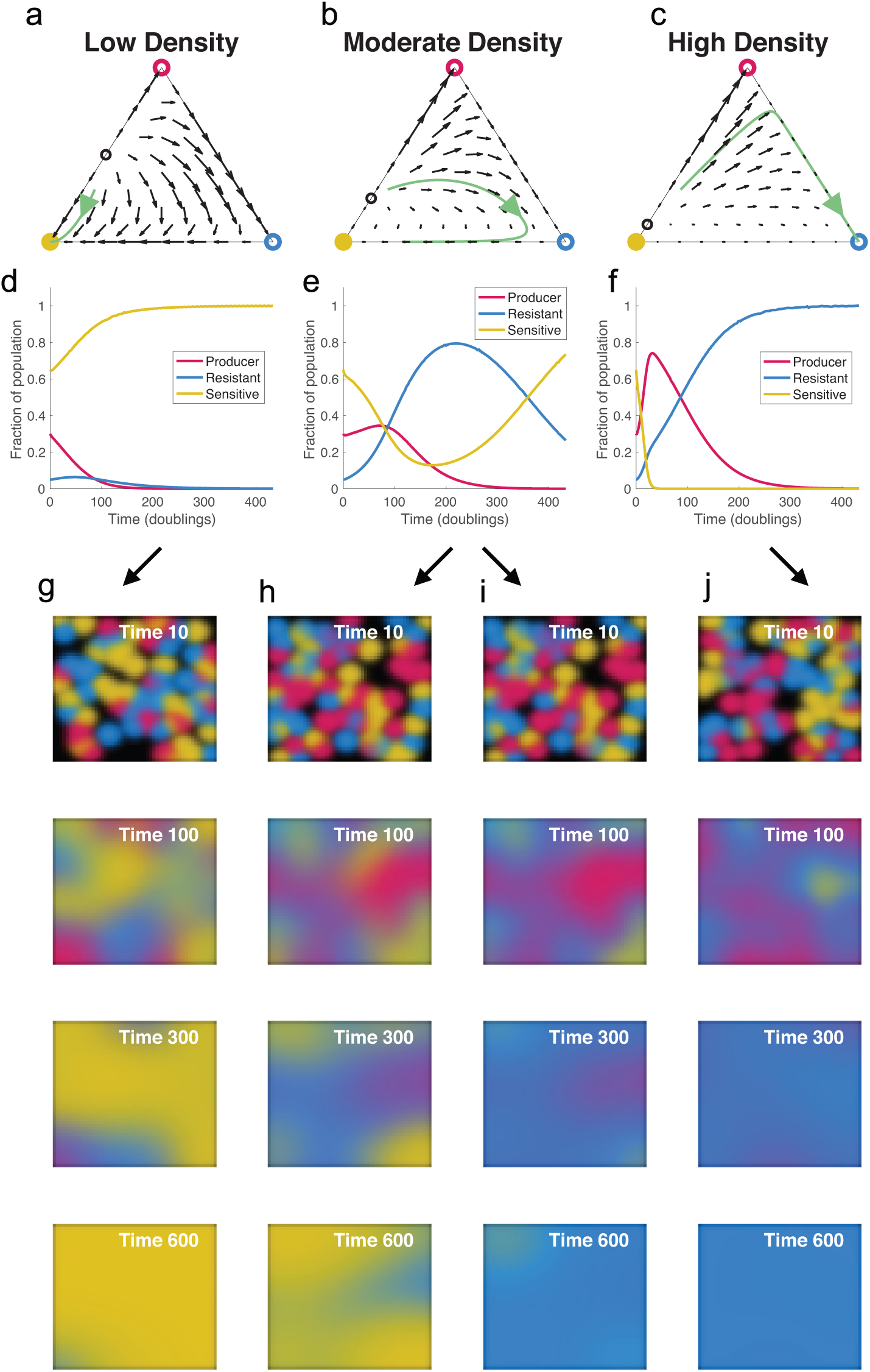
Reaction-diffusion model reveals local and spatial dynamics of T6SS Producer, Resistant and Sensitive genotypes. (**a-c**) Ternary plots showing vector fields and sample trajectories for different values of the density parameter *g* (low density: *g* = 0.1, moderate density: *g* = 0.2, and high density: *g* = 0.5). Each point in the triangle represents a different community composition, with the distance to each edge representing the proportion of each genotype in the population. Open and closed circles at the vertices denote the unstable and stable steady states, respectively. (**d-f**) Progression of community composition for the trajectories shown in (**a-c**). At moderate and high densities, there is alternation of the dominant genotype. (**g-j**) Simulation of the system including spatial structure, where cells can move throughout a 2D surface. For moderate densities, in (**h-i**), the dominant genotype at the end of the simulation depends heavily on the initial distribution of the genotypes across the surface.

To understand the dynamics of a mixed population of Producer, Resistant and Sensitive genotypes, we have so far simulated the system in a well-mixed environment, where interactions are homogeneous across all population constituents. The Producer-Sensitive-Resistant T6SS competition then resembles a Rock-Paper-Scissors game with non-transitive fitness relationships, which has been explored extensively via modeling and experiments ^37^. The model assumption of well-mixed interactions may often be violated in realistic settings, however, including the gut microbiota environment. Spatial constraints limit which cells interact with each other, qualitatively altering the population dynamics of antagonistic (or cooperative) interactions. With this consideration in mind, we next turned to new variations of our model that include spatial constraints on interaction.

We added spatial constraint to our model by assuming that cells can move through a surface, occupying new territory wherever nutrients are still available. We model this movement as diffusion without any directional bias, allowing cells to move freely from higher to lower density areas, resulting in a reaction-diffusion system that tracks the density of each genotype across a 2D space. Provided that the diffusion of cells through the surface is slow enough to retain some spatial structure, we now have regions dominated by each genotype, with interactions happening mostly at the interfaces of these regions (Fig. 5g-j). As a result, the progression of the competition depends on the initial distribution of each genotype across the surface, particularly for moderate colony densities, which are not easily dominated by any single genotype. For instance, the survival of Sensitive cells depends on the existence of pockets shielded from contact with Producer cells, facilitating a takeover once the Resistant genotype replaces the Producer (Fig. 5h-i). Even a minimalistic consideration of spatial constraint on interactions among the Producer, Resistant, and Sensitive T6SS strategies thus begins to recapitulate the retention of sensitive and resistant genotypes akin to what we observed in mouse experiments in which these strategy types were introduced together into the gut environment.

### Individual-based biofilm simulation of T6SS competition

The spatial distributions of species within the microbiome of animal guts, and how they might change through time, are not well known because they are extraordinarily difficult to measure with live microscopy. Outstanding efforts to date have localized different microbiome members to different regions of the intestinal mucosa in addition to the luminal environment of fixed samples of intestinal tissue, or for fluorescently labeled strains in a few live imaging examples ^11,38,39^. Many of these reports have documented communities of multiple strains or species in or on the intestinal mucosa in conditions that resemble biofilm formation, and these surface-bound communities likely undergo local growth combined with different kinds of dispersal. The resulting spatial constraints on movement, localized growth based on gradients of nutrient availability, and dispersal regimes that periodically cull and mix population structure, together are likely to affect how different T6SS genotypes encounter each other, and therefore the relative costs and benefits of Producer, Resistant, and Sensitive strategies ^40–45^.

With these considerations in mind, we implemented simulations of surface-dwelling populations of Producer, Resistant, and Sensitive strains growing together in biofilm-like conditions. Our simulation framework for this system extends from a large family of biofilm models, capturing nutrient diffusion from surrounding the cell populations, local nutrient consumption and growth, and cell-cell shoving as adjacent cell clusters expand or contract (Fig. 6a, Methods) ^46–50^. At any given location, the extent of initial local mixing governs Producers’ access to and ability to kill Sensitive cells, with greater mixing favoring T6SS- producers more and more strongly. As noted above, this dynamic has been described in extensive detail previously as part of the Rock-Paper-Scissors framework ^37,46,51–55^. Rather than recapitulating this work here, we aimed to target the specific questions of how dispersal and re-colonization patterns influence the population dynamics of the three classes of T6SS strains studied experimentally. In separate simulations, we implemented two broad classes of disturbance dynamics, one where the strain composition at a given location was used to seed subsequent locations in a repeating cycle (‘re-colonization’, Fig. 6b), and another in which, within single simulation spaces, randomly sized areas of the substrate were periodically cleared of cells to allow for regrowth (‘sloughing’, Fig. 6c). These two regimes represent disturbance events at relatively large (‘re-colonization’) and smaller (‘sloughing’) spatial scales that could occur as a result of different patterns of mucosal clearance along the length of the intestinal tract.

**Fig. 6.**
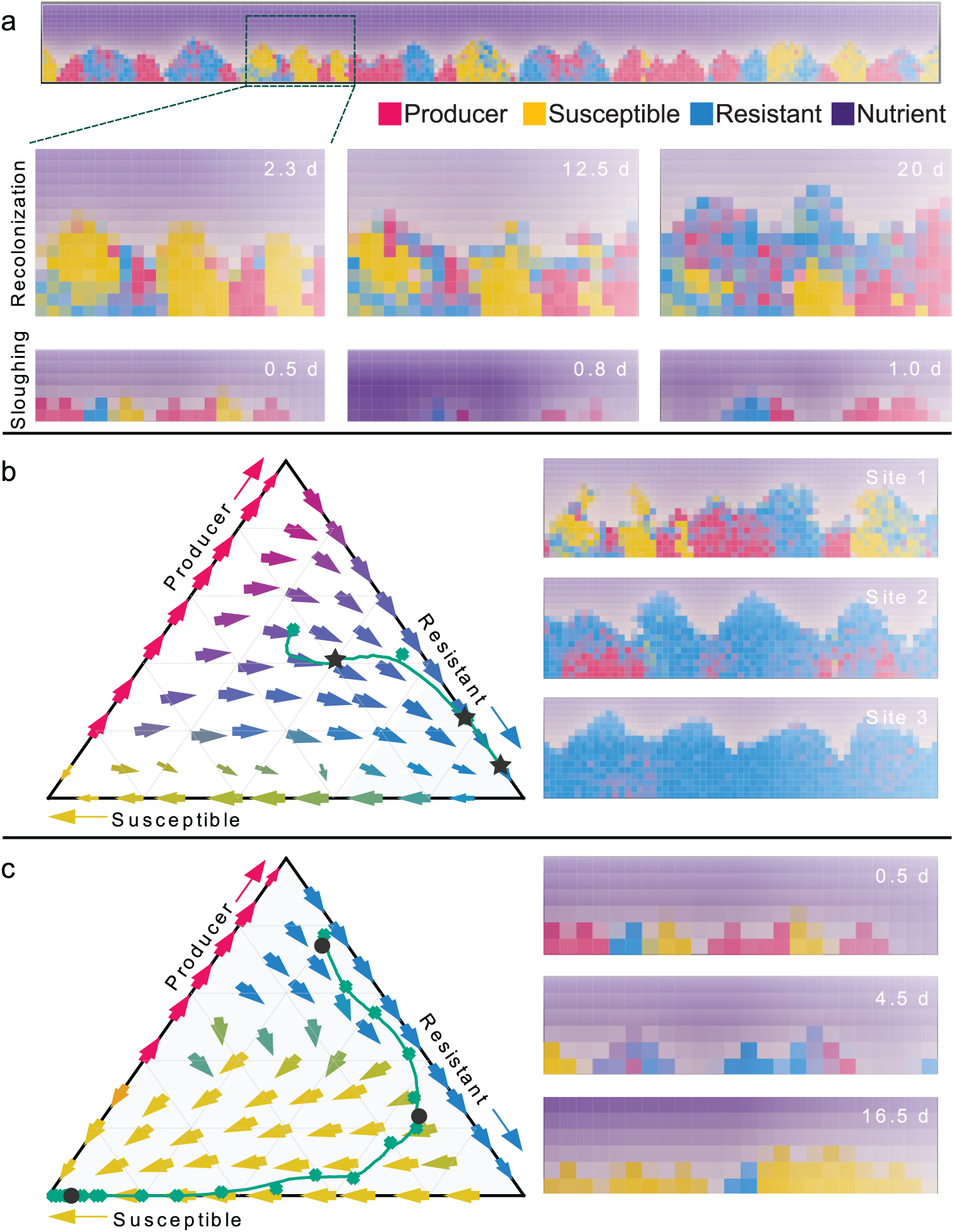
Individual-based biofilm simulations suggest dispersal-recolonization regimes can dictate selective loss of T6SS. (**a**) An example of the biofilm simulation space containing cell groups of T6SS Producer (red), Sensitive (yellow/orange), and Resistant cells (blue). Below the full simulation space, partial frames are shown for the recolonization regime in which cells dispersed from the biofilm surface upstream can re-attach to the biofilm surface downstream, promoting admixture between different strains/species. We also implemented a sloughing regime in which the full biofilm system was periodically cleared and the surface instantaneously recolonized with a sub-sample of the population at the time point just prior to disturbance. (**b**) In the recolonization regime, simulations tended toward mixed regimes containing mostly T6SS Resistant cells, which most closely resembled the mouse experimental outcomes. We note that the pure-Resistant population state is predicted to be invadable in the long term by T6SS Sensitive cells if the Producer strain has been eliminated. (**c**) The sloughing regime, in which biofilm clearance was stronger, tended to generate isolated clonal clusters of cells of each strain along the basal surface. This regime tended more strongly to favor T6SS Sensitive cells, because they remained out of contact with T6SS Producers and had the highest growth rate of the three strain types.

We ran simulations with 1:1:1 inoculation of Producer, Resistant, and Sensitive strains in each of these two disturbance regimes, and found that the re-colonization regime tended toward predominance of the Resistance strain (Fig. 6b). Large-scale disturbance that re-seeds locations with mixtures of the three strains tend to eliminate the susceptible strain, which is continuously re-mixed with the T6SS Producer strains each re-colonization cycle. Consistent contact between Producers and Sensitive cells ensures that the competitive benefit of T6SS production is maintained and offsets the cost of T6SS. After elimination of Sensitive cells, as noted above in our dynamics systems modeling, the Resistant strain displaces the Producer strain because it does not pay the cost of T6SS production. The system tends toward a Resistant-only fixed point (ternary plot in Fig. 6b), but this point is unstable to invasion of Sensitive cells, which can outgrow Resistant. In the sloughing disturbance simulations, on the other hand, particularly if sloughing events are frequent enough that clonal cell groups spend little time in direct contact, the Sensitive strain is predicted to dominate the system and resist invasion by the Producer and Resistant strains, both of which it can out-grow (Fig. 6c). The reason for this shift is that sufficiently fast local disturbance can reduce or eliminate the direct contact of Producer and Sensitive cells, which are rarely killed by T6SS and out-grow the Producer (and Resistant) strains as they pay neither the cost of T6SS nor the immunity factors. Overall, these results suggest that over short to intermediate time scales, which may be what is best represented by our experiments, the Resistant strategy performs best by outgrowing Producer cells, which benefit from killing whatever Sensitive cells they are able to contact. Higher local disturbance and cell group-regrowth increasingly favors the Sensitive strain, which can avoid contact with the Producer and out-grow the Resistant variant. We speculate that this type of micro-environmental turnover may contribute to the fixation of strains that have lost the T6SS machinery on long time scales within host gut microbial communities. The models offer the overall suggestion that intestinal mucosa disturbance and recolonization dynamics may be a critical driver of T6SS population and evolutionary dynamics.

## DISCUSSION

The T6SS is a widespread pathway important for interbacterial antagonism in many taxa. Despite abundant evidence of its prevalence and function in the human gut microbiome, we show here that strains of *B. fragilis* often lose the pathway. Our findings establish that community composition and environmental factors likely modulate the relative costs and benefits of producing the *B. fragilis* T6SS in the human gut. Our work underscores the utility of phylogenomic and metagenomic analysis for the generation of testable hypotheses regarding the mechanisms that underlie the evolution and ecology of the gut microbiota ^31^. While it is common practice to assess the importance of genetic pathways to microorganisms using *in vitro* growth analysis, our experiments indicate that the correct physiological context can reveal fitness costs and benefits that are not otherwise apparent.

Importantly, the gut is not the only context in which T6SS loss has been inferred. For example, *Vibrio cholerae* strains have evolved *de novo* silencing of T6S gene expression in some conditions, or inactivating mutations in T6SS core genes that is also observed for some isolates of *Pseudomonas aeruginosa* derived from chronic lung infections ^56–59^. These observations suggest that some aspects of the evolutionary population dynamics revealed by our studies of *B. fragilis* may also apply to settings outside of the human gut.

Despite the patterns of GA3 loss and the experimental cost of production that we observe in the mouse gut, within most sequenced human gut microbiomes *B. fragilis* retains an intact GA3 system, suggesting that the advantage provided by producing the system is sufficiently large that it outweighs the cost. It is therefore notable that we uncovered a unique association in adults between the abundance of GA3+ *B. fragilis* and two clades of *Bacteroides*, including several highly prevalent and abundant species. *B. fragilis*, among *Bacteroides* species, is well-known for its uniquely close association with the gastrointestinal mucosa, and our results support an *in situ* lifestyle consistent with the importance of contact dependent processes like the T6SS for the natural history of this organism ^60^. Our findings also suggest that cohabitation with certain specific *Bacteroides* species can lead to stronger contact-dependent selective pressure on *B. fragilis* than for other closely-related species. It is striking that the exact clades identified include species that exhibit unusually high prevalence of the GA2 T6SS, which is typically absent from *B. fragilis* genomes ^22^. The association of GA3 with these clades may reflect intra-*Bacteroides* antagonism mediated by distinct T6SS subtypes. Further differences in environmental conditions that cannot be assessed through single-timepoint metagenomic sequencing analysis might facilitate the longer-term evolutionary success of *B. fragilis* GA3 producer strains, including altered mucosal turnover due to diet or genetic factors, or transient ecological disturbances such as antibiotic treatment.

Our phylogenetic analysis revealed that, although GA3 loss is common across the *B. fragilis* phylogeny, lineages lacking GA3 are not evolutionarily successful over long timescales. *Bacteroides* species are well known to be vertically transmitted from mother to infant during birth ^61–63^. The question arises, therefore, if *B. fragilis* loses the T6SS over time in some microbiomes, what is its fate upon vertical transmission during birth to the highly competitive and dynamic ecosystem of the infant microbiome? *B. fragilis* strain replacement has been observed in infants, so these T6SS inactive strains could be replaced by more competitive strains acquired horizontally, thereby providing one potential explanation for the lack of evolutionary persistence of strains lacking GA3 ^24^. An alternate possibility is that, in adults, small reservoirs of *B. fragilis* with functional intact T6SS could remain at levels that are undetectable by our metagenomics analysis but could be sufficient for reseeding the infant gut and upon doing so might rapidly outcompete strains with inactive T6SS.

In keeping with the extensive array of social evolution theory of contact-dependent antagonism, our modeling approaches all indicate that the evolutionary stability of T6SS will depend on the extent of competition for limited resources on local spatial scales, and the extent of intermixing among T6SS Producer, Sensitive, and Resistant cells. For biofilm contexts like those that likely occur in association with the gut mucosa ^11^, our individual-based simulation modeling indicates that under regimes of high disturbance the Producer strain is eliminated rapidly. This outcome may help to explain how T6SS production is selectively eliminated in some host microbiomes, depending on mucosal or epithelial turnover and replenishment regimes that might vary across healthy individuals or in disease states involving altered homeostasis. Future studies should examine the inter-generational population dynamics of *B. fragilis* and the corresponding requirement for costly mechanisms like the T6SS during vertical transmission and establishment in the neonatal intestine. Increasing efforts to resolve the dynamic structure of microbial communities in and on the intestinal mucosa will also be vital to resolving the mechanisms underlying ecological and evolutionary dynamics of key interaction phenotypes including T6SS activity.

## METHODS

### Bacterial culture conditions

*Bacteroides* strains were grown on supplemented brain heart infusion medium (BHIS) plates or liquid cultures supplemented with 1mg/ml of vitamin K3 (Acros Organics/Fisher Scientific, Geel, Belgium) when not specified (Basic 2008)(Lim, 2017) *Bacteroides* strains were incubated under anaerobic conditions (10% CO2, 10% H2, 80% N2) within a 37°C Whitley A55 anaerobic chamber (Don Whitley Scientific, Victoria Works, UK). All BHIS plates contain 60 mg/ml of gentamycin sulfate when not specified. *E. coli* was grown in Luria broth (LB) for liquid or solid agar culture and incubated at 37°C. Antibiotics were added to media as needed: carbenicillin 150 mg/ml, erythromycin 12.5 mg/ml, tetracycline 6 mg/ml, and 2-Deoxy-5-fluorouridine (FudR) 200 mg/ml.

### Plasmid creation

The plasmid pLBG13-Δ*tssBC-clpV* was created by amplifying a right and left homology region flanking the genes *tssBC-clpV* (BF9343_1940-1942) from *B. fragilis* NCTC 9343 genomic DNA template using the primers described in Supplementary Table 4, and Phusion high-fidelity DNA polymerase (New England Biolabs (NEB), Ipswich, MA). Homology arms were cloned via Gibson Assembly into pLBG13 digested with BamHI-HF and SalI-HF restriction enzymes (NEB) using NEBuilder HiFi DNA assembly master mix (NEB). Plasmid sequence was confirmed via Sanger sequencing (Molecular Biology Shared Resources, Dartmouth College, Hanover, NH).

### Genetic manipulation

Integration of pNBU2 plasmids into *att* sites were performed through mating between *Escherichia coli* S17-1 λ *pir* containing the specific pNBU2 plasmid (see Supplementary Table 4) as previously described with some modifications ^64^. For mating, a volume of 25 mL of *E. coli* was incubated in LB to late exponential phase (OD6oo=0.5) on an orbital shaker, and a volume of 3 mL of BHIS media inoculated with *B*. *fragilis* was incubated to exponential phase under anaerobic conditions. Each culture was mixed, pelleted, and washed before plating at high density on non-selective agar media and incubated for 16h at 37°C in an aerobic incubator. The cells were resuspended in 1 mL of BHIS and plates on appropriately selected plates for *B. fragilis*. The insertions were verified by PCR (see Supplementary Table 4). Att1 insertions were used in all cases.

For the generation of in-frame chromosomal deletions using pExchange-*tdk* plasmid and its derivatives, mating was performed as described above using E. coli S17-1 λ *pir* as the conjugal donor strain. After mating, merodiploid integrants were selected on BHIS plates supplemented with erythromycin, streaked to BHIS non-selective media, then resuspended and plated on BHIS media supplemented with FudR for counterselection ^65^. The different mutants were confirmed by PCR. For the generation of mutants using pLBG13 and its derivatives, mating was performed as described previously except for the use of BHIS media for *B. fragilis* ^66^.

For the creation of the strain *B. fragilis* Δ*tdk*Δ*tssBC-clpV,* the mutation in *tssBC-clpV* genes was generated using pLBG13-Δ*tssBC-clpV* in the parental strain. For the creation of the strain *B. fragilis* NCTC9343 Δ*tdk*ΔEIΔ*tssBC-clpV*, the mutation in *tssBC-clpV* genes was generated using pLBG13-Δ*tssBC-clpV* in the strain *B. fragilis* NCTC9343 Δ*tdk*Δ*btei1*, and then the mutation in the gene *btei2* was inserted using pExchange_*tdk*_BF9343_1928-26 in the strain *B. fragilis* NCTC9343 Δ*tdk*Δ*btei1*Δ*tssBC-clpV*.

### Genomic and phylogenetic analysis of *B. fragilis*

We obtained *B. fragilis* genomes from NCBI in November 2021 and ran prokka to identify genes ^67^. We ran blastn to identify homologs of the *B. fragilis* MetaPhlAn3 marker genes against the nucleotide sequences of the predicted genes for each genome ^68^. We used the best hit that had a match of at least 90% of the gene length and a 90% sequence identity. We discarded genomes in which we could identify fewer than 80% of the marker genes as they were likely poor-quality assemblies. We aligned each gene separately using mafft and then concatenated the alignments together for tree construction ^69^. We used RaxML-NG with the GTRGAMMAI model to construct a maximum likelihood tree ^70^. We identified the presence or absence of GA3 by using blastn to find homologs of 13 GA3 structural genes in the *B. fragilis* genomes. We counted homologs with 90% sequence identity over 50% of the length of the gene; the relaxed length criterion was meant to include cases where poor assembly leads to partial gene identification which is of particular concern in regions like the T6SS locus where repetitive sequences are common. Verification of inferred GA3-negative genomes was performed via examination of GA3 loci using Geneious. We reconstructed ancestral states using the ancestral character estimation function (ace) in the R library ape, with a model that estimates different rates for gains and losses (ARD model) ^71^. From those results we identified gains and losses as branches where the presence or absence of GA3 changed. Finally, to calculate the variability in our estimate of GA3 gains and losses, we generated 1000 bootstrapped replicate trees in RaxML and calculated the number of gains and losses for each ^70^.

To assess the significance of the observation that GA3 loss events are close to the tips we used a simulation approach. For each inferred loss event, we calculated the median distance from the branch node where the loss occurred to every descendant tip to quantify how close the loss event was the tips of the phylogenetic tree and determined the median of these distance-to-tip values across all loss events resulting in a single parameter for the entire tree. We then used simulations to assess whether this distance was smaller than expected by chance. We simulated the evolution of GA3 gain and loss events on our phylogenetic tree using a simple two-parameter model: a constant rate of gains and losses per unit length of the tree. These two parameters were inferred using the ace() method in the R library ape used for ancestral reconstruction. We simulated gains and losses of GA3 using a two state markov model as implemented by the rTraitDisc() function in ape. We calculated the same parameter over 1000 simulated tree and compared the observed value to this background distribution.

### Metagenomic studies

We detected the GA3 T6SS in adult metagenomic samples from the Human Microbiome Project (HMP) and infant samples from DIABIMMUNE using similar methodology to what has been previously published ^6,24,28,72^. We compared the number of reads that mapped to the GA3 structural genes to the number of reads that mapped to *B. fragilis* marker genes. The *B. fragilis* marker genes are a subset of the MetaPhlAn markers that are found in all *B. fragilis* reference genomes ^24,68^. Reads were mapped to HMP samples using bowtie2 (parameter -a -N 1) and we counted reads that had at least 95% sequence identity, a quality score of at least 20 and that did not have multiple matches ^73^. We only analyzed samples with at least 100 read counts to *B. fragilis* marker genes to avoid statistical problems associated with low counts. In each sample, we calculated the expected number of reads that mapped to the GA3 T6SS structural genes based on the relative lengths of the two sets of marker genes, and if the actual number was less than 10% of this value we considered it to be GA3-. Species and genus abundance values were calculated using MetaPhlAn v3.0.7 ^68^. Loading vectors were calculated using the code in biplot.pcoa function in the ape package ^71^. PERMANOVA was calculated using the adonis2 function in the vegan package ^74^.

### PhILR analysis

We used the phylogenetic partitioning based isometric log ratio transform (PhILR) package to identify taxa associated with the *B. fragilis* GA3 T6SS in human shotgun metagenomic sequence data ^30^. Briefly, PhILR takes relative microbiome abundance data and applies an isometric log ratio transformation in phylogenetic context to calculate the balance between both descendants on each node of the phylogenetic tree. For this analysis, we used *B. fragilis*-containing microbiome samples that we had defined previously. We supplied the PhILR algorithm with species relative abundance data and a species-level phylogenetic tree (mpa_v30_CHOCOPhlAn_201901_species_tree.nwk) derived from MetaPhlAn3. Because we had previously found an association with *Bacteroides* and furthermore to reduce the burden of multiple testing, we focused our analysis on the phylogeny of *Bacteroides* species. We compared the PhILR abundance values for every node on the phylogenetic tree between GA3 positive and GA3 negative samples using a Wilcoxon rank sum test and corrected for multiple tests using a false discovery rate of 0.05.

### Broth culture growth assays

Growth curves were conducted in defined minimal media with different carbon sources as previously described ^75^. For polysaccharides, we used Orafti®Synergy1 inulin (Beneo, Mannheim, Germany), glycogen (MiliporeSigma, Saint-Louis, MO), and potato starch (MiliporeSigma) at a final concentration of 0.5%. For the other carbon sources (D-glucose (Acros organics), D-fructose (RPI), D-galactose (Acros organics), D-xylose (Alfa Aesar, Haverhill, MA), sucrose (RPI), we used a final concentration of 27mM.

Bacterial strains were streaked on BHIS non-selected plates and incubated under anaerobic conditions for two days. Three colonies per strain were inoculated in 3ml of BHIS media. The cultures were incubated for 20h in anaerobic conditions and back diluted 1:10 in defined minimal media with the appropriate carbon source. The cultures were incubated for 2-3h in anaerobic conditions. The cultures were then diluted in 2ml of the same media at OD600=0.05. Each dilution was used to fill eight wells on a 96-well plate with a volume of 200μl. Media alone was used to fill eight wells on a 96-well plate with a volume of 200μl for the blank. All empty wells were filled with water to limit condensation. The 96-wells plate was covered with Breathe-Easy® gas-permeable membrane (Diversified Biotech, Dedham, MA). The growth was measured every hour in a Stratus (plate reader Cerillo, Charlottesville, VA), and the absorbance was measured at 600nm. The plate reader was incubated in the anaerobic condition described above.

### Solid agar growth assays

Solid growth curves were conducted in defined minimal media with the same carbon sources previously mentioned ^75^. Mucin (MiliporeSigma) was also tested at a 0.5% concentration for the solid growth curve. A 1.5% agar concentration was added to the media to solidify it.

The wild-type strain *att*::pNBU2-*ErmGb* and the mutants ΔBF_1940-1942 *att*::pNBU2-BC14-*tetQ* were streaked on BHIS-selected plates with either erythromycin or tetracycline and incubated in anaerobic conditions for two days. Three colonies per strain were inoculated in 3ml of defined minimal media with the appropriate carbon source. The cultures were incubated for 20h in anaerobic conditions. A volume of 2 ml of overnight cultures was washed once in PBS and resuspended in PBS. The strains were diluted at OD600=3.00. For the monocultures, the volume of each replicate was mixed to an equal volume of PBS. For the mixed, both strains were mixed in equal volume to a 1:1 ratio. For both conditions, monocultures and mixes, multiple spots were inoculated on defined minimal media with the tested carbon sources. The initial inoculum of the monocultures and mixes are quantified by calculating the colony-forming unit (CFU) of each strain on both BHIS-selected plates with erythromycin and tetracycline. At different time points (5h, 10h, 24h, 29h, 34h, 48h), the plates were removed from the anaerobic chamber. For each replicate, one spot was cut from the agar and transferred in an Eppendorf with 1ml PBS and a glass bead (4.5mm). The tubes were vortexed for 30 seconds and the suspensions were used to calculate the CFU of each strain on both BHIS-selected plates with erythromycin and tetracycline.

### Interbacterial competition assays

The competitions were performed as described previously with some modifications ^26^. *B. fragilis* NCTC 9343 Producer strains were co-cultured at a 10 to 1 ratio (OD600 6.0 to 0.6) with either resistant or susceptible strains on BHIS agar containing 5% horse blood (Colorado Serum Co.). Producer and Susceptible strains were resistant to either erythromycin or tetracycline. Competitions were incubated for 16 to 18 hours under anaerobic conditions at 37°C. Competition outcomes were quantified by enumerating colony-forming units (CFU) for each strain by plating on BHIS agar media supplemented with appropriate concentrations of either erythromycin or tetracycline.

### Animal studies

All animal experiments were performed using protocols approved by the Dartmouth College Institutional Animal Care and Use Committee (protocol number 00002231). Mice used were Swiss Webster (SW) females received at 8-10 weeks of age (health status murine pathogen-free, MPF) purchased from Taconic Laboratories (Hudson, NY, USA). The mice were housed under SPF conditions in the Dartmouth Center for Comparative Medicine and Research. Prior to colonization assays, mice were treated with a three-course antibiotic regiment to eliminate the endogenous microbiota and allow engraftment, as described previously ^76^. Fecal samples were collected prior to oral gavage. Each group of mice was composed of eight animals housed in pairs and experiments were performed with at least two independent biological replicates using mice from different litters. No statistical tests were used to pre-determine number of mice in experiments.

Bacterial strains used in mouse experiments possessed unique barcode sequences (see Supplemental Table 4) ^77^. To prepare bacteria for gavage into mice, the strains were streaked on non-selective BHIS agar plates and incubated overnight under anaerobic conditions at 37°C. The bacterial cells were then resuspended in phosphate buffered saline (PBS) and were washed by pelleting before strains were mixed at a 1:1 or 1:1:1 ratio with a final viable cell count of 10^8^-10^9^ cfu/ml in a 15% glycerol solution in PBS and stored at -80°C until gavage. Immediately prior to gavage, aliquots of strain mixtures were thawed, and the mice received 100-200 μl of the thawed preparations.

Fecal samples were collected at different intervals in a 3- or 6-months period after gavage. Half of the pellets were frozen at -80°C and the other half was resuspended in glycerol 15% and was frozen at - 80°C. At necropsy, samples were taken along the length of the gut for some mouse groups. For the fecal pellet, a fraction was used for isolating colonies on BHIS-tetracycline selected plates.

For the fecal pellets conserved in the -80°C without glycerol from the different timepoint, the genomic DNA (gDNA) of the pellets was extracted using Dneasy Blood and Tissue kit (Qiagen, Hilden, Germany), and 5 ng of each DNA extraction was used for qPCR. Quantification of each strain was performed by qPCR as described previously using CFX96 Detection System (Bio-Rad) and SsoAdvanced master mix (Bio-Rad, Hercules, CA) with the primers described in Supplemental Table 4 ^77^. The standard curve was prepared with gDNA from pure cultures of the barcoded strains (range 2ng, 0.2ng, 0.02ng, 0.002ng, 0.0002ng) and was used to calculate the relative abundance of each strain.

### Assessment of T6SS function and genomic sequencing in evolved isolates

Single colonies were isolated from fresh fecal pellets at days 187 or 180 of the two 6 months mouse experiments from select mice. Briefly, pellets were resuspended in LB, supernatant removed from debris, and serially diluted. Each dilution was plated on BHIS agar media supplemented with tetracycline and incubated under anaerobic conditions. Single colonies isolated in this manner were re-streaked to ensure purity. Producer and Resistant evolved clones were identified via PCR amplification of the BC01 and BC14 barcodes. Following identification of each colony, T6SS function was assessed via ELISA protocol as described previously by Weber *et al.* with some modifications ^78^. Briefly, isolated were inoculated in BHIS into a 5mL culture tube or wells of a 96-well plate with parental strains as controls and incubated overnight under anaerobic conditions. The stationary-phase cultures were then back-diluted 1:10 in fresh BHIS broth and incubated 4h under anaerobic conditions. OD600 was assessed using a Synergy Neo2 plate reader (Biotek, Winooski, VT). The cultures were centrifuged 10 minutes at 5,000xg and supernatant were centrifuged a second time in the same condition. The binding step was carried out overnight at 4°C prior to incubation for 90 minutes at room temperature in blocking solution (5% bovine serum album (BSA) in phosphate buffered saline). The antibodies solutions was composed of 2.5% BSA in PBS-T. Primary anti-TssD polyclonal antibodies were gifted by Harris D. Bernstein ^34^. Secondary antibodies donkey anti-rabbit IgG HPR conjugated (Biolegend, San Diego, CA) was used with a 1:7500 dilution. The reaction was stopped with 2M of sulfuric acid and the OD450 was measured for quantification. Five biological replicates are represented in Figure S6.

For whole genome sequencing, the evolved clones were streaked on BHIS blood agar plates and incubate overnight under anaerobic conditions. Cells were then resuspended in PBS and genomic DNA was extracted using the Dneasy Blood and Tissue DNA extraction kit (QIAGEN). Genomic DNA was also extracted from the Ross lab parental isolate of *B. fragilis* NCTC 9343 as a reference. Whole genome sequencing was performed by the Microbial Genome Sequencing Center (MIGS, Pittsburgh, PA) using Illumina NextSeq 2000 (Illumina, San Diego, CA). Reads were aligned to the *B. fragilis* NCTC 9343 reference genome and variants detected using breseq version 0.35.4 ^79^.

### 16S rRNA gene amplicon sequencing of mouse endogenous microbiota

For taxonomic profiling of the endogenous microbiota during the mouse experiment, 16S rRNA gene sequencing was performed on DNA extractions from fecal pellets. DNA extractions were the same as used for qPCR quantification of *B. fragilis* strain abundances. Extracted DNA samples were analyzed by first amplifying the V4-5 hypervariable regions of the 16S ribosomal RNA (rRNA) gene. A first-round PCR amplified the target region using universal primers 16S-515FB-PCR1_NX (5’- TCGTCGGCAGCGTCAGATGTGTATAAGAGACAGGTGYCAGCMGCCGCGGTAA-3’) and 16S-926R-PCR1_NX (5’-GTCTCGTGGGCTCGGAGATGTGTATAAGAGACAGCCGYCAATTYMTTTRAGTTT-3’). Using the Illumina Nextera Index kit, a second round of PCR amplification was then performed to introduce the indexes and adaptors according to the manufacturer’s recommendations. Amplified PCR reactions were subsequently prepared and sequenced at the University of New Hampshire Hubbard Center for Genomics Sequencing Core Facility using an Ilumina HiSeq2500 with custom sequencing primers added to the reagent cartridge and sequenced 2 × 250bp ^80^. The sequence data were deposited in the NCBI SRA under BioProject ID ----------------. Resulting community sequences were processed using R as outlined below. Complete analysis code is available in supplemental materials and via GitHub: https://github.com/mcclur51e/TypeSixDisadvantage.

For ASV taxonomy assignment, raw (adaptor-trimmed) sequences were processed using the standard DADA2 pipeline ^81^. Sequences were filtered using standard filtering parameters: 0 Ns, truncated quality score less than 2, primers trimmed, and maximum expected error rate of 2. Forward and reverse sequences were then merged, and chimeras removed to reach a dataset containing 13,070 unique ASVs. Singleton ASVs were trimmed from the dataset to reach a dataset containing 4,492 unique ASVs. ASVs were assigned taxonomy by comparing to the SILVA SSU 132 dataset ^82^. Any ASVs that were not identified to the genus level and were considered significant in later analysis were compared to the RDP dataset for taxonomic assignment to the lowest possible taxonomic level ^83^.

Three types of negative controls were prepared and sequenced. 1) Reagent controls were prepared by performing the DNA extraction procedure using the same reagents without any sample. 2) Negative-PCR controls were prepared by performing first round PCR amplification using molecular biology grade water. 3) Negative submission controls contained molecular biology grade water with no first round PCR. Two types of positive control were prepared and sequenced. 1) Amplification of a ZymoBIOMICS Microbial Community DNA Standard (Zymo Research, Irvine, CA U.S.A.) and 2) amplification of DNA extracted from a ZymoBIOMICS Microbial Community Standard (Zymo Research, Irvine, CA U.S.A.). Contaminating ASVs were identified through the use of the decontam package ^84^. After removal of contaminants identified by decontam, the dataset consisted of 4,315 ASVs. Samples were further trimmed to remove reads from Animals, Eukaryotes, and Chloroplasts and resulted in a dataset consisting of 4,222 bacterial ASVs. Samples contained 25 – 942,687 reads, with a median = 37,616 (median = 52 for negative controls). The trimmed dataset was then transferred to Phyloseq for manipulation ^85^. Images were prepared using ggplot2 and cowplot.

### Population dynamics simulations

To model the dynamics of producer (*p*), resistant (*r*) and sensitive (*s*) subpopulations, we consider competition for resources between the different genotypes, with a fitness cost for resistance and production of T6SS, as well as contact killing of the sensitive genotype by the producer. We model the growth of the different genotypes using coupled differential equations for logistic growth while competing for resources, with *K* denoting the carrying capacity and *γ* denoting the maximum growth rate of the sensitive genotype^36^. The coupling of the equations implements direct competition for resources, with the abundance of one genotype being limited by the abundance of the other. Since all genotypes are clonal except for the presence of T6SS production or resistance, we consider that they occupy the same ecological niche and therefore have the same carrying capacity. We model the fitness costs of production of T6SS *α_P_* and resistance *α_R_* as a reduction in maximum growth, with 0 < *α_P,R_* < 1 and *α_p_* > *α_R_*. Killing of the sensitive genotype happens with a certain probability, whenever a producer cell encounters a sensitive cell. In a well-mixed environment, killing is therefore proportional to both the sensitive and producer populations, with a rate parameter *δ*. We have then the following system of equations:

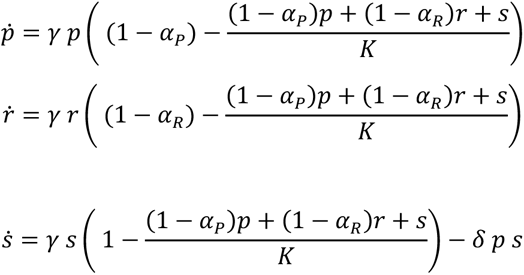

Next, we scale the cell populations by the carrying capacity (*p*’ = *p*/*K*, *r*’ = *r*/*K*, *s*’ = *s*/*K*) and scale time by the maximum growth rate (*t*^’^ = *t γ*) to arrive at a system of non-dimensional equations (we drop the prime hereafter for simplicity):

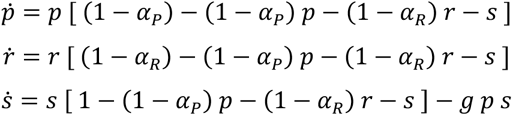

The parameter *g* = *δ K*/*γ* is the killing rate per contact scaled by the carrying capacity and standardized to the non-dimensional time measure 1/*γ* (we further discuss this parameter in the main text). We simulate this system using MatLab ODE solver *ode113* to integrate the system of equations over time.

We model movement of the different genotypes across a 2D surface by adding a diffusion term, with the same scaled diffusion constant for all genotypes, arriving at a reaction-diffusion model consisting of a system of partial differential equations:

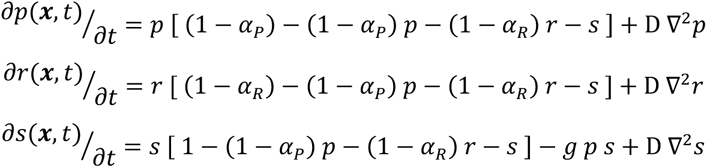

We then simulate the model for an initial condition where the genotypes are randomly scattered across the surface. We use the Crank–Nicolson method ^86^, choosing time steps and distance increments such that Δ*t* < 0.5 Δ*h*^2^ / *D* to guarantee the stability of the solution.

#### Model Parameters Table

|  |  |  |
| --- | --- | --- |
| Killing rate scaled by carrying capacity | $g$ | variable |
| Cost of production | $\alpha_p$ | 2% |
| Cost of resistance | $\alpha_R$ | 3% |
| Diffusion constant of genotypes across surface | $D$ | 0.0001 |

### Biofilm simulations

Here we use the agent-based simulation framework previously described ^48^. Briefly, a simulation space is defined as a two- or three-dimensional space where nutrients diffuse from the top of the system down toward bacteria attached in the substratum. As the bacteria take up these nutrients, they grow in mass and divide. The bacteria in the system are assumed to be in small well-mixed boxes or grid-nodes, 3 μm in side-length, and when the volume of the bacteria in a node is larger than the volume allows, they overflow into a neighboring box. The system is assumed to be under flow, causing an erosive force removing individuals from the surface of the biofilm. The erosion can cause a mass of biofilm to be effectively cut off from the substratum, and all biomass that isn’t connected to the substratum is removed^87^.

As a result of nutrient consumption on the biofilm’s advancing front, nutrient gradients are created with high nutrient availability in the outer cell layers and lower nutrient availability with increasing depth into the biofilm interior. Cells near the liquid interface grow with the maximum growth rate, while cells deeper in the biofilm interior grow relatively slowly. Fluid flow is modeled implicitly; following prior literature, we allow the biofilm to erode along its outer front at a rate proportional to the square of the distance from the basal substratum (described in detail in ^49^).

The steps of the simulation per iteration include:

1. diffusion of nutrients
2. bacterial growth and division
3. erosion and detachment of biomass
4. sloughing (if applicable)
5. recolonization
6. Type VI secretion system activity
7. biofilm relaxation (overflowing grid nodes)

The simulations in this study use a T6SS where each individual with a T6S effector checks for individuals in the same grid node. Randomly selecting one, if that individual lacks the corresponding immunity gene and the system successfully fires then it is killed and removed. A successful ‘fire’ occurs at a probability determined from the rate parameter in the table below. Each immunity and effector has a cost which is removed from the biomass at the time of bacterial growth.

Mixing methods introduced for this work include recolonization and sloughing. Recolonization is implemented for each individual that is removed from the system via detachment or erosion, retain the individual in a buffer at some rate (probability, given fixed *dt*). In subsequent iterations, re-introduce the retained individuals at another rate. These rates are the retention and recolonization rates, respectively. To re-introduce a particle, they are simply added to a random location on the surface of the biofilm. Sloughing is implemented as a simple 3 step process. First, calculate the frequency of the three strains of bacteria in the system. Next, remove all particles from the system. Finally, re-inoculate the system at the previously calculated frequency.

The relative order of most steps is biologically motivated as in previous work ^48^, with the exception of the T6SS, which is in order for computational efficiency. In testing, this selection had no effect (SI).

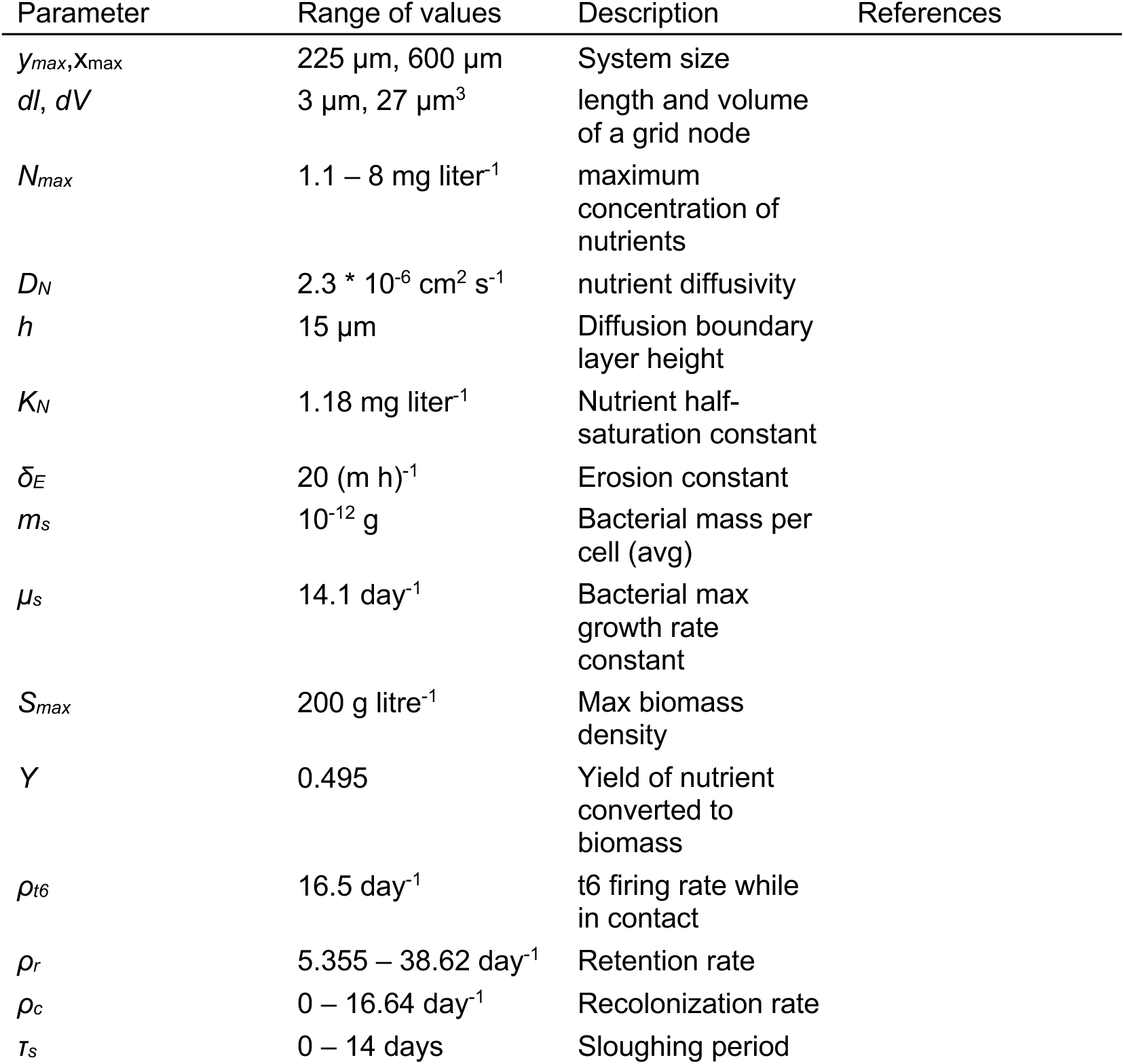

## Acknowledgments

We thank the members of Dartmouth College’s Joint Evolutionary Microbiology Meeting (JEMM) and Microbiology and Microbial Pathogenesis (M2P2) communities and Joseph Mougous for feedback on this work; Dr. Nicholas Pudlo and Dr. Eric Martens for technical advice; BENEO for providing Synergy inulin; Dr. Harris Bernstein for anti-TssD antibodies; Dr. Andrew Goodman for providing *Bacteroides fragilis* strains and plasmids. This work was supported by start-up funding from Dartmouth College Geisel School of Medicine and NIH grants R00GM129874 and R35GM142685 to B.D.R.; NSF grant IOS 2017879, Simons Foundation award number 826672, and HFSP award number RGY0077/2020 to CDN; NIH P20 GM130454 to DS; and the Dartmouth College BioMT Core funded by P20GM113132.

## Extended Data Figure Legends

**Extended Data Fig. 1.**
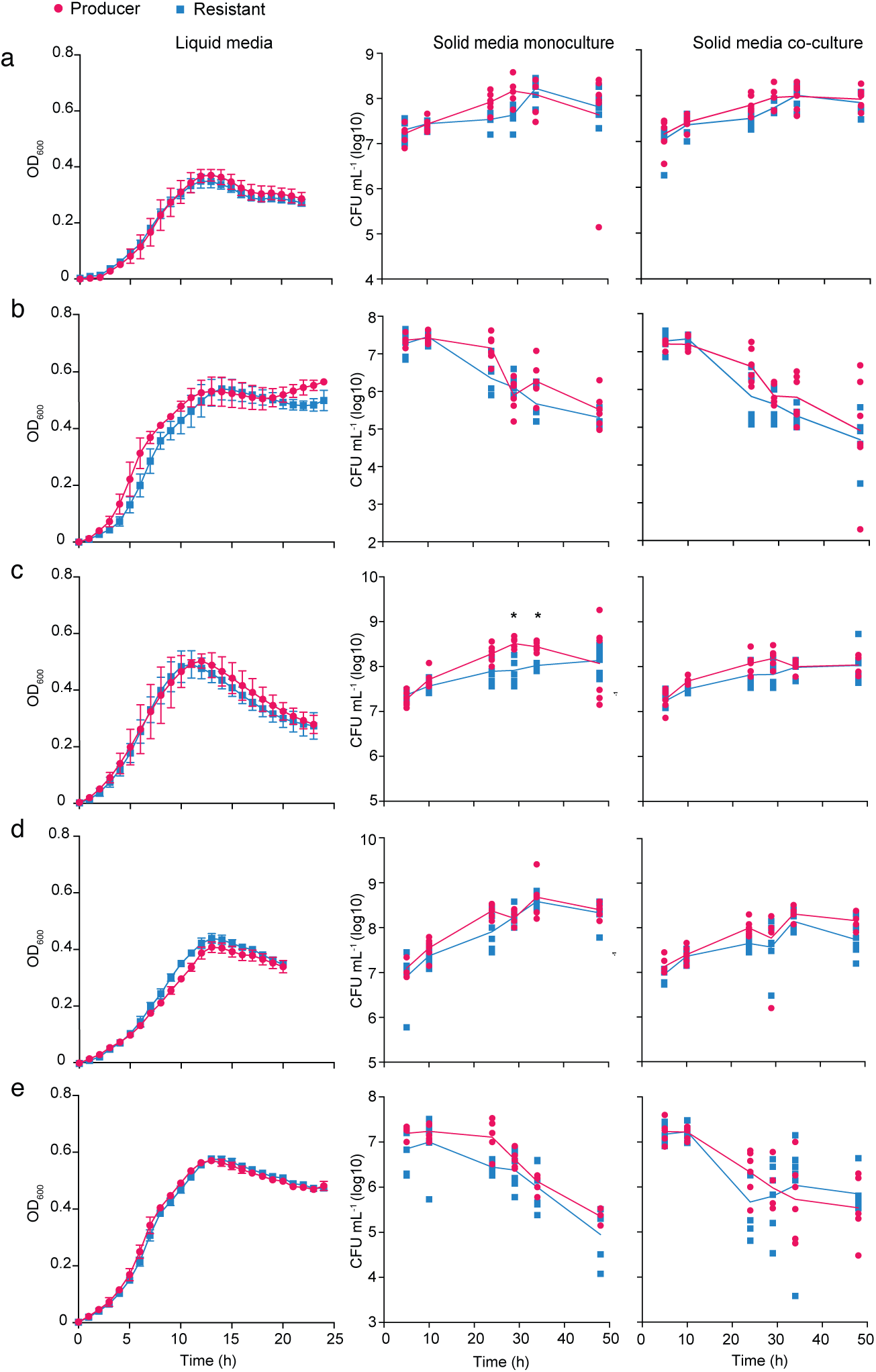
Abrogation of T6SS activity does not provide a fitness benefit to *B. fragilis* under laboratory conditions growing on mono and di-saccharide as carbon source. Growth curves and solid growth in mono and co-culture for the Producer and Resistant strains subjected to growth on different sole carbon sources (mono and di-saccharide) in defined minimal broth or solid agar media, including D-glucose (**a**), D-fructose (**b**), D-galactose (**c**), D-xylose (**d**), and sucrose (**e**). For solid agar growth co-culture, producer and resistant strains were mixed at a 1:1 ratio by OD600, inoculated in high-density spots before being harvested for enumeration of colony forming units by plating on BHIS supplemented with either erythromycin or tetracycline. Mann-Whitney tests at each timepoint indicated no significant difference between strains. For liquid experiments, the results are from three biological replicates that each represent the mean of eight technical replicates. For solid growth, results are from six biological replicates from two independent experiments. Data shown represent mean values ±SD.

**Extended Data Fig. 2.**
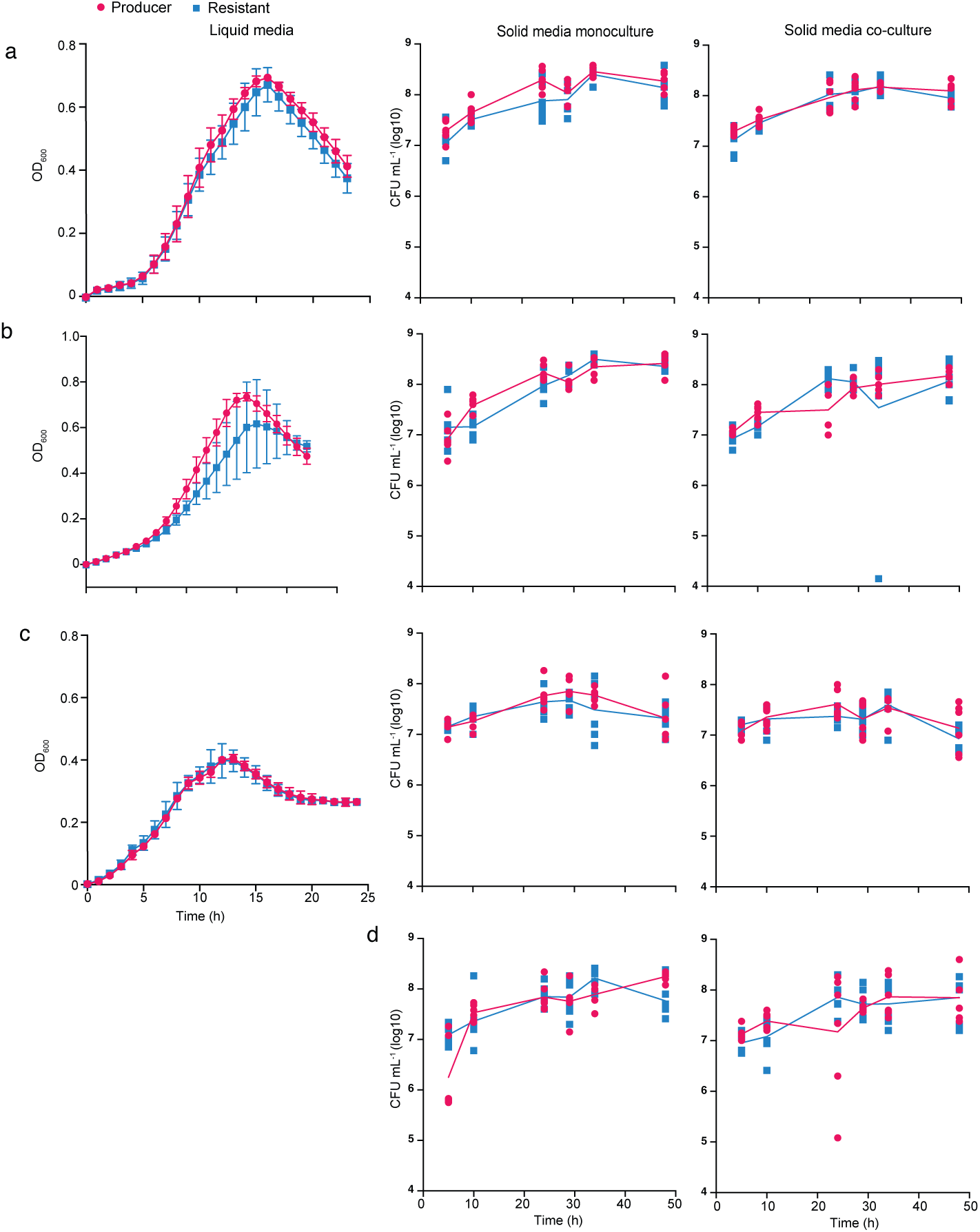
Abrogation of T6SS activity does not provide a fitness benefit to *B. fragilis* under laboratory conditions growing on polysaccharides as carbon source. The growth of the Producer and Resistant strains in defined minimal media with different carbon sources (polysaccharide) is shown in liquid and solid (mono and co-culture) growth in the different panels. For solid growth, *B. fragilis* Producer strain with Erm^R^ and Resistant strain with Tet^R^ were grown on defined minimal media with different carbon sources separately (monoculture) and together in a 1:1 ratio (co-culture). Each strain was quantified via calculating colony-forming units on BHIS agar plates supplemented with erythromycin or tetracycline. The bacteria grew in defined minimal media supplemented with starch potato (**a**), glycogen (**b**), ©Synergy Inulin (**c**), and mucin (**d**). Mann-Whitney test showed no difference between the different strains at each timepoint. For liquid experiments, the results are from three biological replicates that each represent the mean of eight technical replicates. For solid growth, results are from six biological replicates from two independent experiments. Data shown represent mean values ±SD.

## Supplemental Figure Legends

**Fig. S1.**
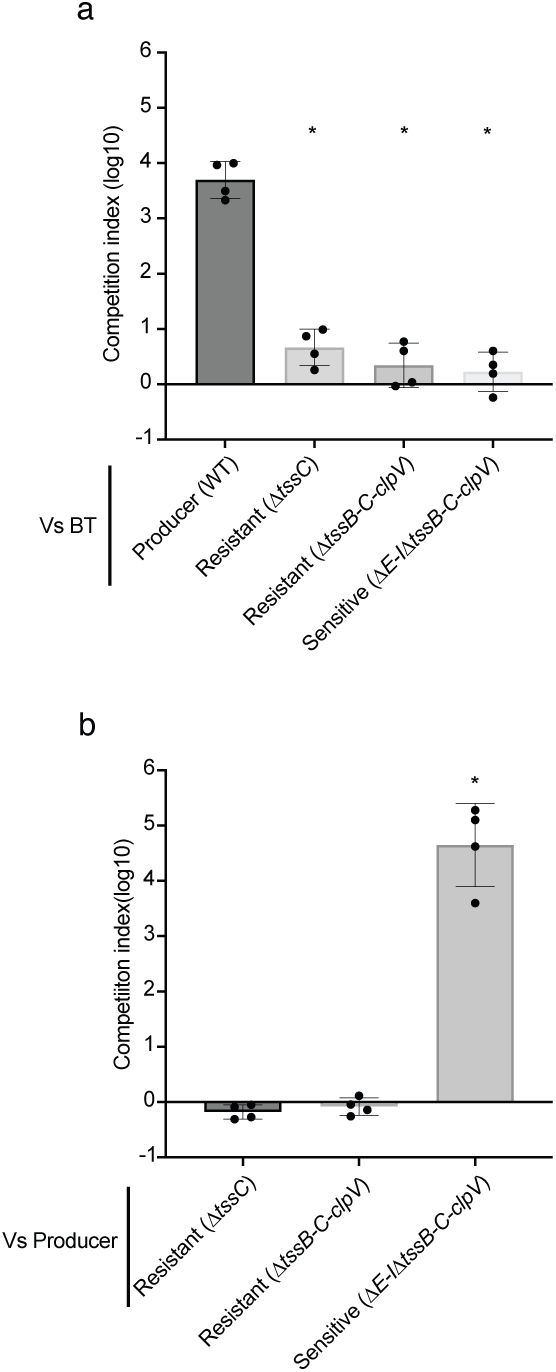
Competition tests of the T6SS producer strain and the T6SS resistant strain. Strains used for the growth experiments were evaluated for their T6SS-dependent competitive phenotype through (**a**) interspecific co-culture competition against a natural Sensitive strain *(Bacteroides thetaiotaomicron*) (BT) or (**b**) tested for sensitivity to T6SS-dependent intoxication through intraspecific competition against the WT Producer strain. Competitive index values for co-culture competitions used to evaluate T6SS activity in (**a**) reflect *B. fragilis*:*B. thetaiotaomicron* ratios. Competitive index values for co-culture competitions used to evaluate sensitivity to T6SS activity in (**b**) reflect Resistant:Producer or Sensitive:Producer ratios. Mann-Whitney tests used to compare mean competitive index of Producer (WT) vs BT **(a)** or Resistant(Δ*tssC*) vs Producer across conditions (*P<0.05). Results are from four independent biological replicates each with three technical replicates. Data show the mean values ±SD.

**Fig. S2.**
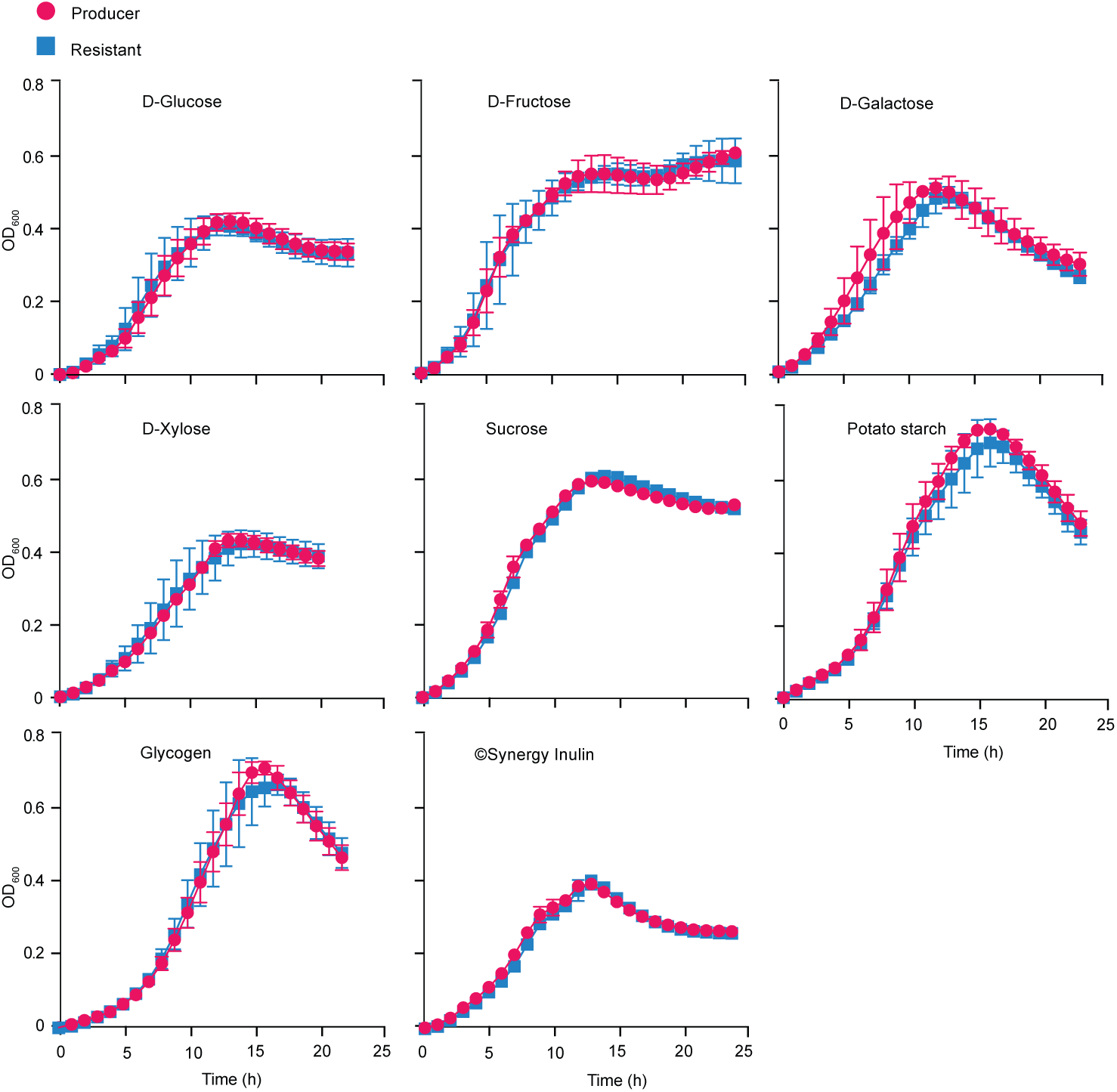
Liquid growth assays involving different mono-, di-, or polysaccharides as carbon sources. The growth of T6SS Producer (WT) and Resistant strains (Δ*tssC*) in defined minimal media supplemented with different carbon sources: D-glucose, D-fructose, D-galactose, D-xylose, sucrose, potato starch, glycogen, and inulin. Mann-Whitney test showed no difference in means between the different strains at each timepoint. For each experiment, the results are from three biological replicates that are the mean of eight technical replicates. Data show the mean ±SD.

**Fig. S3.**
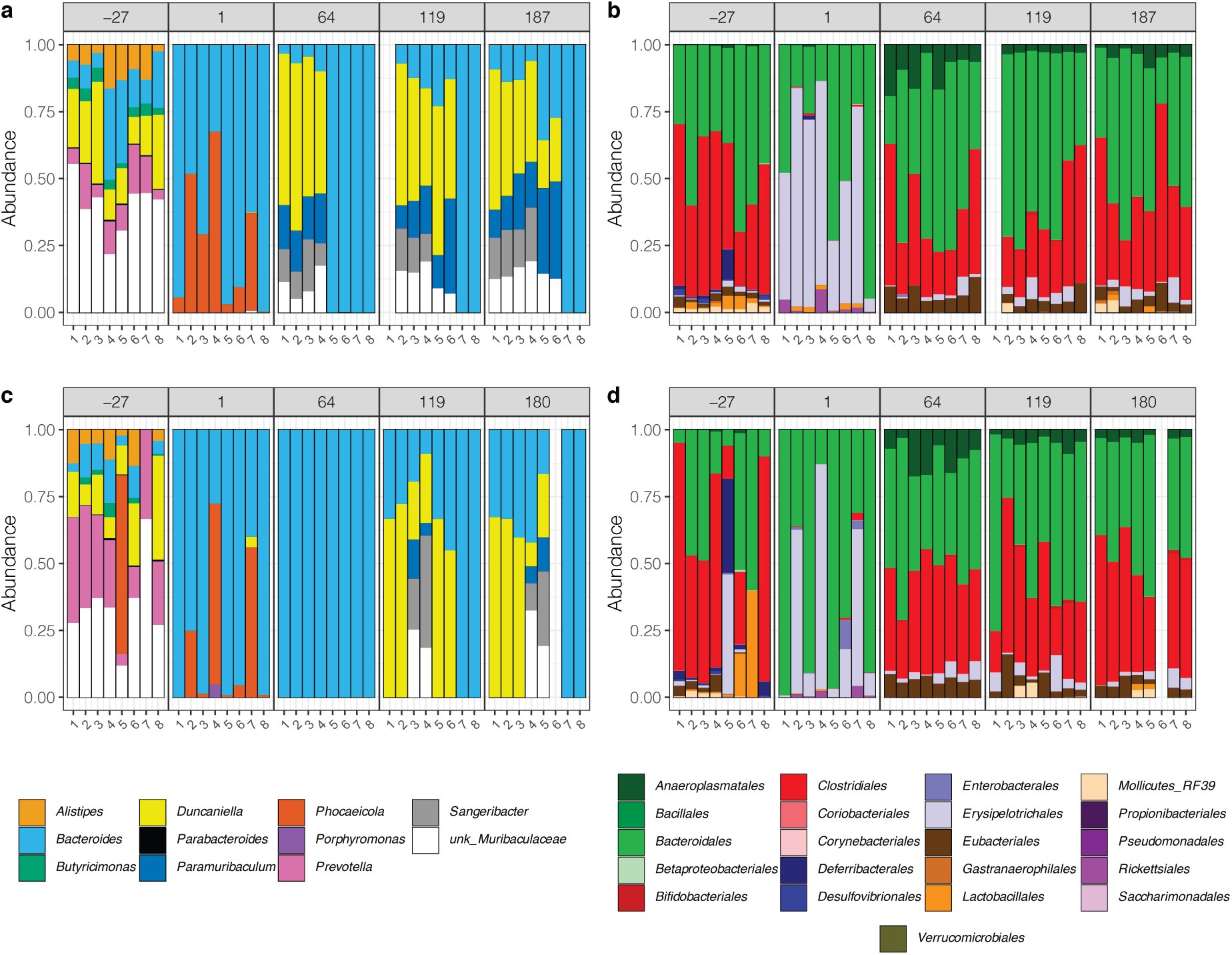
Taxonomic profiling of microbiota from two-strain co-colonization experiments in mice. Taxonomic profiling of the gut microbiota via 16S rRNA amplicon sequencing after 6 months of co-colonization of mice by the Producer strain with a Resistant strain either (**a-b**) lacking *tssC-B-clpV* genes or (**c-d**) a Resistant strain lacking *tssC* only. Taxonomic profiles are shown at the genus level (**a, c**) for taxa within the order Bacteroidales or the order level (**b, d**) for all taxa.

**Fig. S4.**
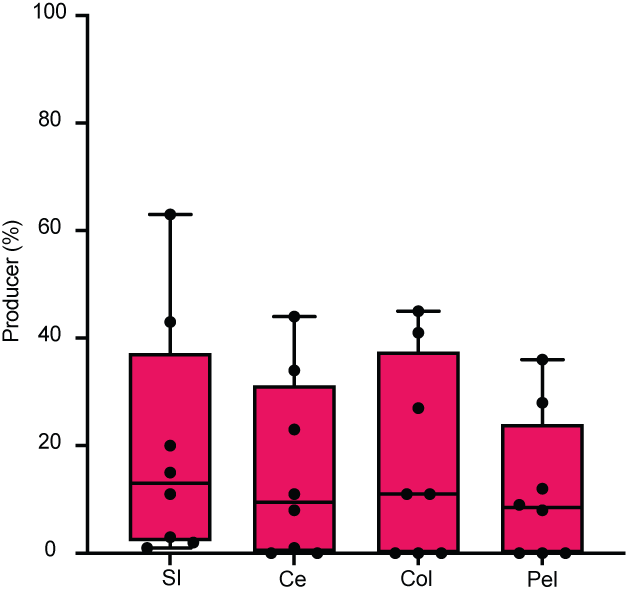
Quantification of strain relative abundance along the gastrointestinal tract from co-colonization of mice. The percent of Producer compared to the total of Producer and Resistant strains (Δ*tssBC-clpV*) detected along the GI tract at day 187 after gavage (SI: Small Intestine; CE:Cecum, Col:Colon, Pel:Pellet), calculated by qPCR targeting unique barcodes. One-way ANOVA analysis showed no difference between mean strain abundances across different intestinal sites (P>0.05).

**Fig. S5.**
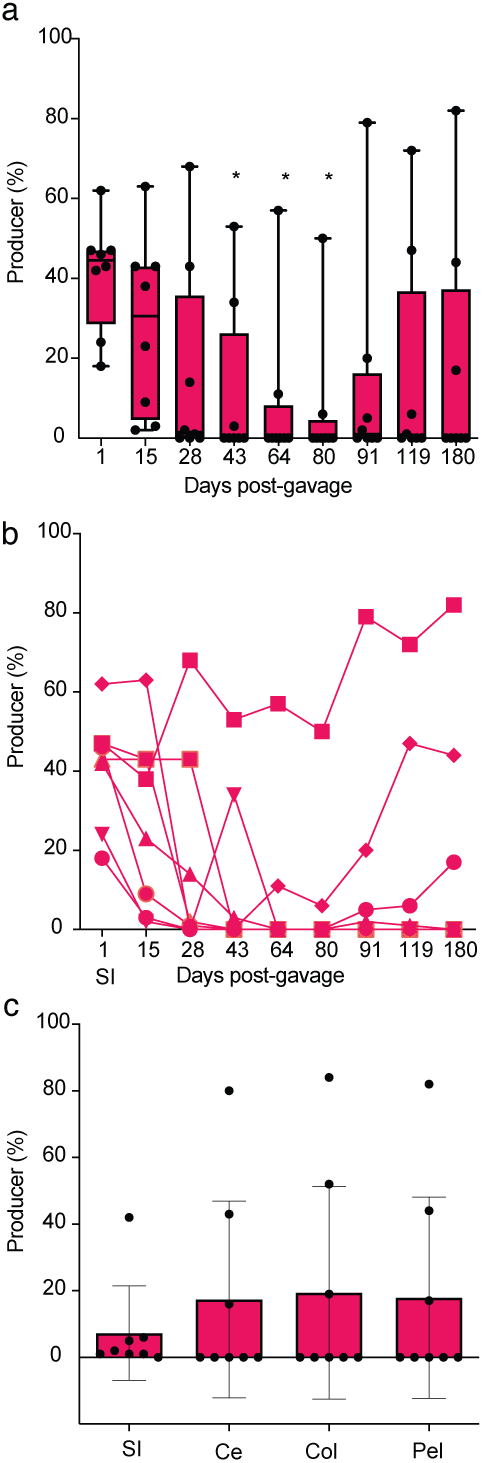
Co-culture of T6SS producer and T6SS resistant strains in the mouse gut. The T6SS producer (WT) and T6SS resistant strain (Δ*tssC)* were inoculated in a 1:1 ratio in mice pre-treated to eradicate the endogenous gut microbiota. **(a)** We followed the two strains progression for 180 days (around 6 months) by sampling the mice pellet multiple times. The T6SS producer percentage is shown with a mean of eight. Kruskal-Wallis test showed the difference between sampling at day 1 (one day after gavage) and all subsequent sampling. *P < 0.05 (**b**) T6SS Producer percentage follow-up of each mouse. (**c**) Percentage of gDNA producers detected along the GI tract at day 180 after gavage (SI: Small Intestine, CE:Cecum, Col:Colon, Pel:Pellet). Kruskal-Wallis showed the absence of difference between the different localization (P>0.05).

**Fig. S6.**
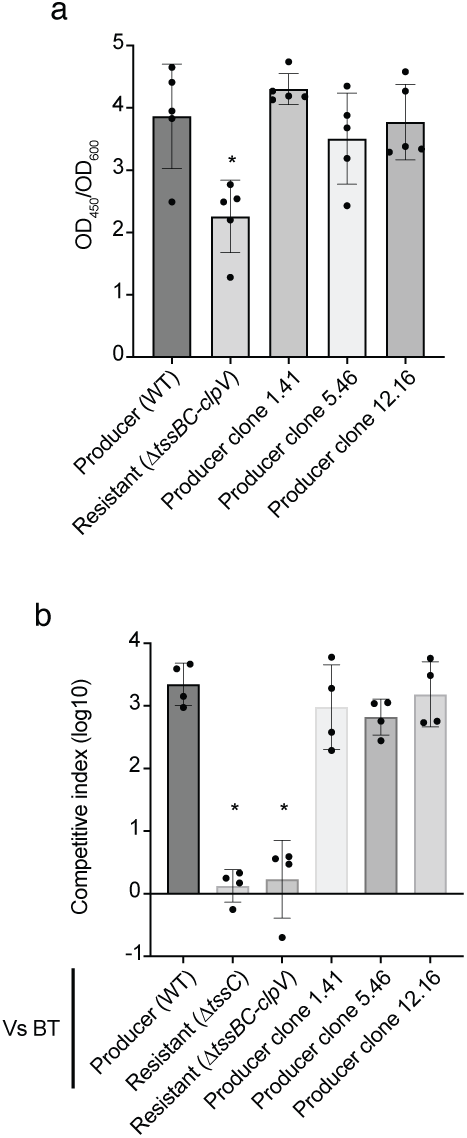
Producer strains do not lose T6SS activity following extended co-colonization of mice. WT clones were isolated from the last fecal pellets of the two six months experiment (Fig.2 and Fig.S5). Screening for the absence of Hcp secretion was performed but none of the clones showed a loss of secretion. (**a**) Three clones were randomly chosen, and detection of secreted Hcp is shown by an ELISA using Hcp-specific antisera. Mann-Whitney test was used to compare parental WT strain to all other strains (*P<0.05). Results are from five biological replicates. (**b**) T6SS activity was further tested for the same isolates used in (**a**) via two-strain co-culture growth assays using *B. thetaiotaomicron* (BT) as a susceptible target strain. Mann-Whitney test compare WT vs BT against all the other conditions (*P<0.05). Data points represent the mean of three technical replicates for each of four biological replicates ±SD.

**Fig. S7.**
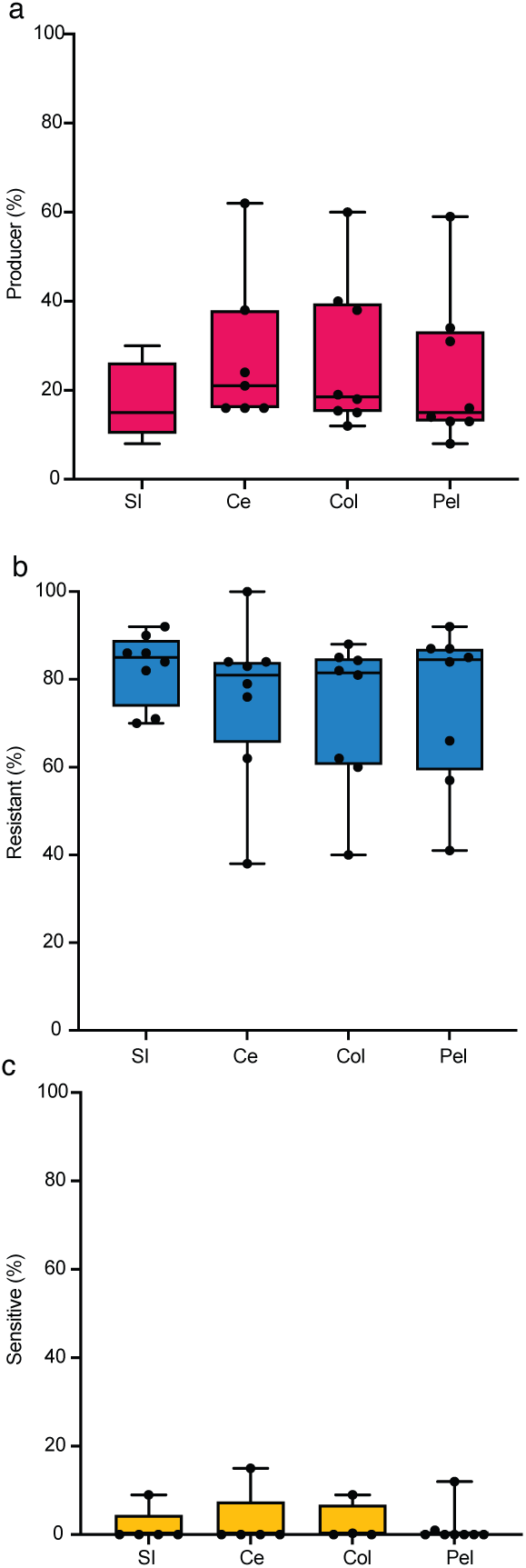
Quantification of strain relative abundance along the gastrointestinal tract from three-strain co-colonization of mice. The T6SS Producer (WT), Resistant strain (Δ*tssBC-clpV)*, and Sensitive strains (Δ*E-I*Δ*tssBC-clpV*) were inoculated in a 1:1:1 ratio in mice pre-treated to eradicate the endogenous gut microbiota. After 88 days, the contents of the small intestine (SI), cecum (CE), colon (Col), were collected upon sacrifice. The mean percent relative abundance of Producer (**a**), Resistant (**b**), and Sensitive (**c**) strain by quantification of barcodes via qPCR is shown for each mouse individually over time. D’Agnostino & Pearson tests were used to determine normal distribution of data and one-way ANOVA (for Resistant strain) and Kruskal-Wallis (for Producer and Sensitive strains) tests indicated that no significant differences between strain composition across intestinal cites.

**Supplementary Table 1.** Classification of adult gut metagenomes by *B. fragilis* GA3 T6SS status and taxonomic relative abundances for species identified to be associated with GA3 via PhILR.

**Supplementary Table 2.** Reference genomes of *B. fragilis* annotated by GA3 status.

**Supplementary Table 3.** Genomic differences identified in *B. fragilis* Producers evolved in mice co-colonized with Resistant strains.

**Supplementary Table 4.** Strains, plasmids, and primers used in this study.

